# Integrative genomic analysis identifies unique immune environments associated with immunotherapy response in diffuse large B cell lymphoma

**DOI:** 10.1101/2024.01.17.576100

**Authors:** Sravya Tumuluru, James K. Godfrey, Alan Cooper, Jovian Yu, Xiufen Chen, Brendan W. MacNabb, Girish Venkataraman, Yuanyuan Zha, Benedikt Pelzer, Joo Song, Gerben Duns, Brian J. Sworder, Christopher Bolen, Elicia Penuel, Ekaterina Postovalova, Nikita Kotlov, Aleksander Bagaev, Nathan Fowler, Sonali M. Smith, Ash A. Alizadeh, Christian Steidl, Justin Kline

## Abstract

Most diffuse large B-cell lymphoma (DLBCL) patients treated with bispecific antibodies (BsAb) or chimeric antigen receptor (CAR) T cells fail to achieve durable treatment responses, underscoring the need for a deeper understanding of mechanisms that regulate the immune environment and response to treatment. Here, an integrative, multi-omic approach was employed to characterize DLBCL immune environments, which effectively segregated DLBCLs into four quadrants – termed DLBCL-immune quadrants (IQ) - defined by cell-of-origin and immune-related gene set expression scores. Recurrent genomic alterations were enriched in each IQ, suggesting that lymphoma cell-intrinsic alterations contribute to orchestrating unique DLBCL immune environments. In relapsed/refractory DLBCL patients, DLBCL-IQ assignment correlated significantly with clinical benefit with the CD20 x CD3 BsAb, mosunetuzumab, but not with CD19-directed CAR T cells. DLBCL-IQ provides a new framework to conceptualize the DLBCL immune landscape and uncovers the differential impact of the endogenous immune environment on outcomes to BsAb and CAR T cell treatment.

## Introduction

Immunotherapies such as chimeric antigen receptor (CAR) T cell therapy and bispecific antibodies (BsAbs) have revolutionized the treatment landscape of relapsed or refractory (r/r) diffuse large B cell lymphoma (DLBCL). For example, CD19-directed CAR T cell therapy induces durable responses in ∼30-40% of patients with r/r DLBCL (1–4). Complete responses are also observed in up to 40% of r/r DLBCL patients who receive CD20 x CD3 BsAbs (5–7). However, in many r/r DLBCL patients, BsAbs and CAR T cell therapy either fail completely or confer only fleeting responses. Collectively, these clinical observations indicate that while immunotherapies can be effective against DLBCL, additional research is needed to understand the cellular and molecular features of the DLBCL environment that underlie immunotherapy responsiveness or resistance to enrich for the subset of patients who will derive maximal benefit and to identify strategies to overcome mechanisms driving intrinsic or acquired resistance.

Numerous studies have demonstrated that a “T cell-inflamed” environment identifies solid cancers against which a spontaneous anti-tumor immune response has been raised and serves as a useful biomarker of response to checkpoint blockade therapy (CBT) (8–11). Transcriptional and immune cell profiling of DLBCL has also revealed subsets enriched in immune cell infiltration and in expression of immune-related genes, which indicates that some DLBCLs exhibit evidence of having stimulated an endogenous anti-lymphoma immune response (12–15). However, the critical factors that shape the DLBCL immune environment are unclear and the impact of the immune environment on promoting or preventing immunotherapy response has yet to be fully defined (16,17). Therefore, in this study, we utilized an integrative approach to 1) broadly characterize the DLBCL immune landscape, 2) provide insight into the contribution of genomic alterations in shaping the DLBCL immune environment, and 3) define the extent to which transcriptionally based immune clustering of DLBCL informs responsiveness to CD20 x CD3 BsAb and CAR T cell immunotherapy. Through these analyses, we uncovered four unique DLBCL immune environments (or immune quadrants - IQs), each enriched for several distinct oncogenic alterations, suggesting that lymphoma cell-intrinsic factors play an important role in orchestrating the immune environment in this disease. For example, we identified loss-of-function (LoF) mutations in *SOCS1* as significantly enriched in DLBCLs with inflamed environments, and that genetic ablation of *Socs1* renders B cells more sensitive to the effects of T cell effector cytokines such as interferon-gamma (IFNγ). Finally, we find that DLBCL-IQ assignment impacts clinical outcomes to BsAb but not CAR T cell treatment, which suggests a differential effect of the endogenous immune environment on DLBCL outcomes to these two immunotherapy treatments.

## Results

### Transcriptomic analysis identifies four immune-related DLBCL clusters

In solid cancers, a “T cell inflamed” gene signature identifies tumors against which a spontaneous anti-tumor immune response has been induced and enriches for a subset of cancers that are particularly vulnerable to immunotherapies, such as CBT (8–11). To determine if DLBCLs could be similarly segregated, we performed bulk transcriptional analyses focused specifically on immunological aspects of the lymphoma environment. Our discovery cohort consisted of transcriptomic data from diagnostic DLBCL biopsies from the National Cancer Institute (NCI, n = 481) (18,19) and an internal dataset from the University of Chicago Medical Center (UCMC, n = 96) (20), comprising a total of 577 cases (*See Methods for details)*.

Gene set variation analysis (GSVA) (21) was used to calculate sample-wise relative enrichment scores for 19 gene sets that fell into 2 functional groups: 1) twelve immune-related signatures that reflect the presence or activation state of immune cell subsets, and 2) seven cell-of-origin (COO) - related signatures **(Supplementary Table 1)**. Immune-related signatures were manually curated from an extensive literature search and contained gene sets related to: IFNγ response (8), type I IFN response (11), CD8^+^ T cell activation (11,22,23) and exhaustion (11), T helper subsets (T_h_1 (22), T_h_2 (22), T follicular helper (T_fh_) (22), and regulatory T cells (T_reg_) (11)), macrophages (11), and dendritic cells (24). As COO segregates DLBCLs into activated B-cell-like (ABC) and germinal center B-cell-like (GCB) subtypes with distinct clinical, transcriptional, and genetic features, we hypothesized that COO would also significantly influence the composition of the DLBCL immune environment. Therefore, we included gene sets derived from a validated gene expression-based COO classifier (25), as well as gene sets regulated by IRF4 (26) and BCL6 (26), transcription factors critical for COO classification. Altogether, 577 unique genes were included in the 19 gene sets selected for GSVA (< 15% overlap between individual gene sets).

GSVA scores for the 577 DLBCLs in the discovery cohort were next subjected to principal component analysis (PCA), which revealed that the majority (69.8%) of variance in gene set expression scores was explained by PC1 (45.38%) and PC2 (24.43%). Unsupervised k-means clustering performed on the PCA-transformed dataset revealed that the optimal number of clusters was four **(Supplementary Figure 1A)**. DLBCLs in our discovery cohort were then assigned to one of four specific clusters described below. Samples from NCI and UCMC datasets were equally represented in each cluster, indicating the absence of significant batch effects **(Supplementary Figure 1B and C)**.

Individual immune-related gene set expression scores were highly correlated, trending in the same direction along PC1 **(Figure 1A)** and made minimal contributions to PC2 **(Supplementary Figure 1D and E)**, indicating that PC1 represented an immune-related axis. In support of this notion, differential gene expression (DGE) analysis of putative “hot” DLBCLs (PC1 > 0) and “cold” DLBCLs (PC1 < 0) revealed upregulation of genes linked to immune cell infiltration and activation (*CD2, CD3D, GZMK, IFNG, CD8A, PRF1*) in the former **(Supplementary Figure 1F)**. ABC COO-related gene sets (ABCDLBCL-1, ABCDLBCL-2, IRF4Up-7) were highly correlated and positively associated with PC2, while GCB COO-related gene sets (GCBDLBCL-1, GCBDLBCL-2, IRF4Dn-1) were negatively associated with PC2, indicating that PC2 represented a COO-related axis **(Figure 1A, Supplementary Figure 1D and E)**. Comparative analysis of the transcriptomes of putative “ABC” DLBCLs (PC2 > 0) and “GCB” DLBCLs (PC2 < 0) identified several genes known to drive COO classification **(Supplementary Figure 1G)**. Orthogonal validation demonstrated that GSVA-defined “ABC” (PC2 > 0) and “GCB” (PC2 < 0) DLBCLs exhibited >95% concordance with COO designations from established COO classifiers, confirming our GSVA was accurately classifying DLBCLs based on COO **(Figure 1B and C)** (15,25). Thus, immune-related and COO-related gene sets are orthogonal and segregate DLBCLs into 4 quadrants based on a distinct distribution of immune-related (PC1) and COO-related (PC2) scores, referred to as DLBCL-immune quadrants (DLBCL-IQ). Specifically, ABC DLBCLs could be transcriptionally defined as “ABC hot” (n = 182, 31.5%, PC1 > 0, PC2 > 0), characterized by high enrichment scores for ABC COO-related and immune-related gene sets, or “ABC cold” (n = 184, 31.9%, PC1 < 0, PC2 > 0). Similarly, GCB DLBCLs could be subdivided into “GCB hot” (n =122, 21.1%, PC1 > 0, PC2 < 0) and “GCB cold” (n = 89, 15.4%, PC1 < 0, PC2 < 0) clusters **(Figure 1D and E)**.

**Figure 1.**
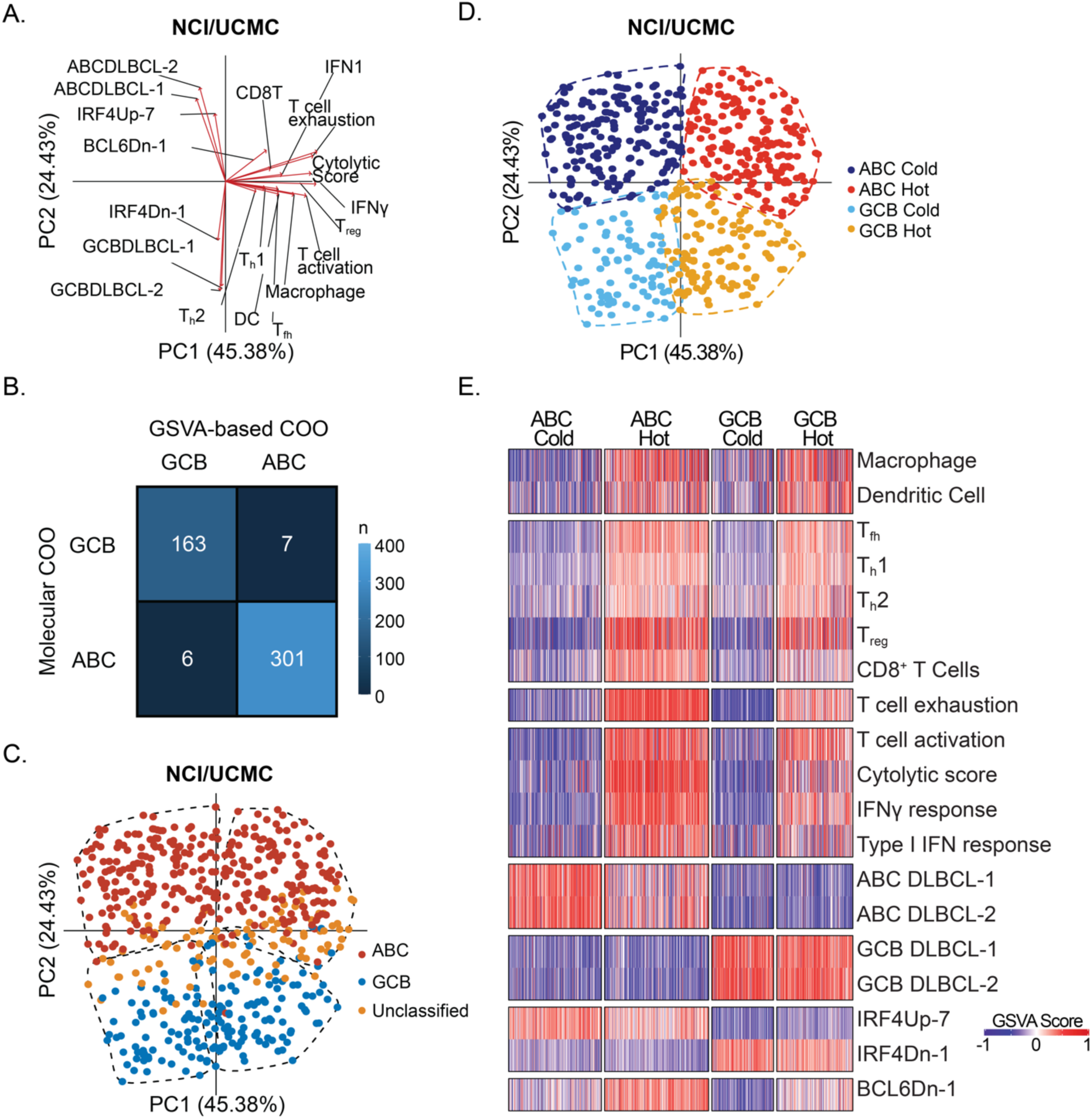
Transcriptomic analysis identifies four unique DLBCL immune environments. **A.** Principal component analysis (PCA) biplot showing the contribution of immune-related gene sets and COO-related gene sets to PC1 and PC2, respectively. **B**. Confusion matrix showing the concordance between GSVA-based COO classification and molecularly defined COO designations for all DLBCLs in the discovery cohort (NCI/UCMC, n = 577). **C.** PCA plot showing concordance between sample-wise GSVA COO scores and molecularly defined COO calls for all DLBCLs in the discovery cohort. Clusters are defined by the dashed line. **D.** PCA plot showing sample-wise GSVA enrichment scores for DLBCLs in the discovery cohort, labeled by immune quadrant (IQ) name (ABC Cold – dark blue, ABC Hot – red, GCB Cold – light blue, GCB Hot – yellow). **E.** Heatmap showing sample-wise GSVA enrichment scores for all 19 gene sets.

When an identical GSVA was performed on an independent DLBCL dataset (British Columbia Cancer, BCC, n = 285) (27,28), a similar clustering pattern emerged **(Supplementary Figure 2A)**, with immune-related gene sets contributing to PC1 and COO-related gene sets contributing to PC2 **(Supplementary Figure 2B)**. Therefore, the NCI/UCMC and BCC datasets were combined to power downstream analyses. In the combined dataset (n = 862), DLBCLs from different sources (NCI/UCMC versus BCC) were similarly distributed among each of the four IQs **(Supplementary Figure 2C and D)** indicating an absence of batch effect. Moreover, the combined dataset was again distributed into four distinct IQs, demonstrating the stability of GSVA-based clustering in classifying DLBCLs by COO and their immune environment **(Supplementary Figure 2E and F)**.

### Validation of transcriptionally defined DLBCL-IQs

Next, to confirm that DLBCL-IQs accurately reflect the actual immune cell composition of specific lymphoma environments, multispectral immunofluorescence (mIF) was performed on a subset of DLBCLs (n = 65, UCMC) for which paired RNA-seq data were available. Staining was performed using fluorescently labeled antibodies against T cell markers (CD4 and CD8), macrophage markers (CD68), DAPI, and B cells (PAX5). Macrophage (CD68^+^) to DLBCL (PAX5^+^) cell ratios were similar across immune clusters **(Supplementary Figure 3A)** and were not significantly correlated with sample-wise hot/cold (PC1) axis scores **(Supplementary Figure 3B)**. However, there were significant differences in T cell infiltration among the four DLBCL- IQs. Representative images of a hot and cold DLBCL are shown in **Figure 2A**. DLBCLs transcriptionally clustered by GSVA as ABC or GCB hot were characterized by significantly higher ratios of CD8^+^ T cells to DLBCL cells when compared to ABC and GCB cold counterparts **(Figure 2B)**. Furthermore, CD8^+^ T cell to DLBCL cell ratios were significantly correlated with sample-wise hot/cold (PC1) axis scores **(Figure 2C)**. The same was true for CD4^+^ T cell to DLBCL cell ratios (**Figure 2D and E**). These data validate the accuracy of GSVA-based clustering in classifying DLBCLs as harboring hot or cold immune environments.

**Figure 2.**
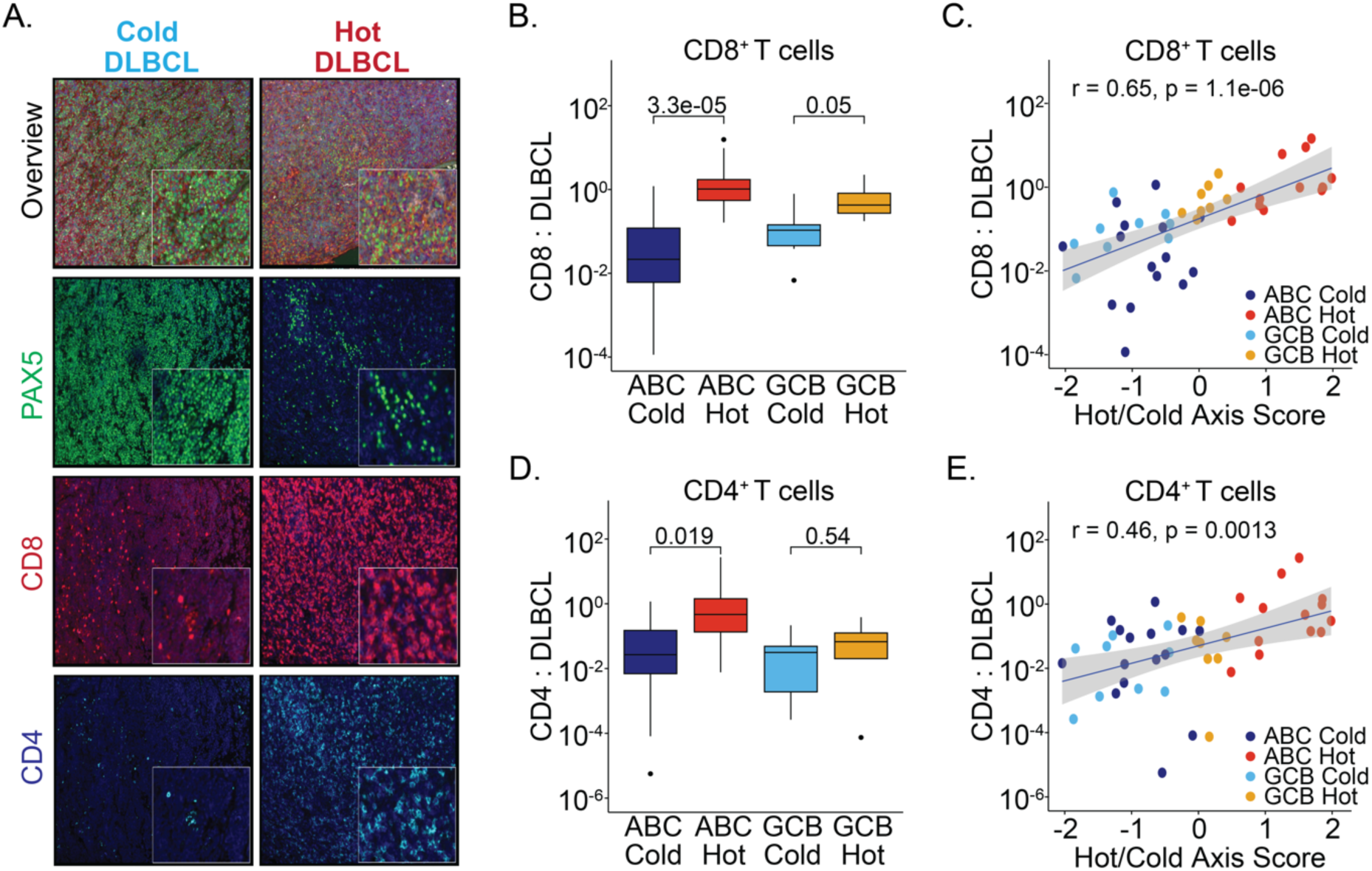
Validation of transcriptionally defined DLBCL-IQs. **A.** Representative multispectral immunofluorescence (mIF) images showing Pax5^+^ lymphoma cells (green), CD8^+^ T cells (red), CD4^+^ T cells (blue) for a cold (left) and hot (right) DLBCL. **B.** Box plot showing average CD8^+^ T cell : DLBCL cell ratios in immune-related clusters (n = 45). **C.** Scatter plot showing correlation of hot/cold axis score (PC1) and CD8^+^ T cell : DLBCL cell ratio (n = 45). **D.** Box plot showing average CD4^+^ T cell : DLBCL cell ratios in immune-related clusters (n = 45). **E.** Scatter plot showing correlation of hot/cold axis score (PC1) and CD4^+^ T cell : DLBCL cell ratio (n = 45).

### Prognostic significance of DLBCL-IQs

Next, the extent to which our DLBCL-IQs correlated with clinical outcomes to initial treatment with R-CHOP was assessed. As expected, DLBCLs assigned to GCB clusters showed improved progression-free survival (PFS) and overall survival (OS) compared to DLBCLs assigned to ABC clusters. However, further stratifying within ABC or GCB clusters according to hot or cold IQ assignment did not impact survival outcomes. **(Supplementary Figure 4A and B)**.

### Concordance of DLBCL-IQs with other immune-related DLBCL subtypes

Recently, DLBCLs have been classified according to the composition of their immune environments. For example, Kotlov and colleagues recently classified DLBCLs into four lymphoma microenvironments (LMEs) based on functional gene expression signatures (F^GES^) that reflected the relative abundance and functional features of cellular constituents (15). When LME cluster annotations were applied to DLBCLs in our combined NCI/UCMC/BCC dataset, we found that DLBCLs assigned to the ABC hot cluster overlapped significantly with an inflammatory microenvironment (LME-IN) **(Supplementary Figure 5A)**. GCB and ABC cold DLBCLs were equally represented within the depleted environment (LME-DE) **(Supplementary Figure 5A)**. Finally, germinal center-like (LME-GC) and mesenchymal (LME-ME) did not overlap significantly with any GSVA-defined immune clusters **(Supplementary Figure 5A)**.

Additionally, Steen et al. have introduced Ecotyper, a molecular classifier that integrates immune cell deconvolution with single-cell RNA sequencing to characterize diverse DLBCL cell states and ecosystems within the DLBCL environment. When comparing the overlap between our DLBCL-IQs and the 9 lymphoma ecotypes (LE), we noted that approximately 65% of DLBCLs in the ABC cold cluster were classified into LE1 and LE2, which are distinguished by the highest expression of ABC COO-related genes and the greatest abundance of B cells, while being deficient in T cell subsets **(Supplementary Figure 5B)**. Similarly, around 55% of GCB cold DLBCLs were assigned to LE8, which is marked by an increased abundance of B cells and the expression of GCB-associated genes **(Supplementary Figure 5B)**. Notably, DLBCLs in the LE8 cluster exhibited significant overlap with DZsig^+^ DLBCLs. Finally, LE4, characterized by enrichment in ABC-related genes and an abundance of CD4^+^ T cells, CD8^+^ T cells, NK cells, and Tregs, predominantly consisted of ABC hot DLBCLs. Minimal overlap was observed between the remaining LEs and our GSVA-based IQs. Collectively, these results suggest LME and Ecotyper capture different aspects of the lymphoma microenvironment and only partially overlap with our DLBCL-IQs.

### Genomic features associated with DLBCL-IQs

Emerging evidence indicates that specific oncogenic alterations and associated transcriptional programs can significantly impact the composition of the tumor immune environment and vulnerability to immunotherapies in solid tumors (29–37). Therefore, we analyzed paired whole exome sequencing (WES) and copy number alterations (CNA) for DLBCLs in the combined cohort to identify genetic features associated with each of our GSVA-based DLBCL-IQs.

In DLBCL, several landmark studies have developed genomic approaches to describe new DLBCL subtypes based on the presence of specific co-occurring genetic alterations, including a probabilistic classifier called LymphGen (18,38,39). However, the association of specific LymphGen clusters with distinct immune environments has not been fully elucidated. LymphGen cluster annotations were available for 754 DLBCLs in our combined dataset. Broadly speaking, LymphGen classifications overlapped imperfectly with our DLBCL-IQs **(Figure 3A)**, with a few notable exceptions. For example, DLBCLs characterized by activating *NOTCH1* mutations (LymphGen N1 subtype), were significantly enriched in the ABC hot cluster, while DLBCLs with *TP53* loss and aneuploidy (LymphGen A53 subtype), were almost exclusively found in the ABC cold cluster. Lastly, DLBCLs with gain-of-function mutations in *EZH2*, *BCL2* translocations, and *MYC* activation (LymphGen EZB-MYC^+^ subtype), were enriched in the GCB cold cluster. The remaining four LymphGen subtypes showed little correlation with specific GSVA-based immune-related clusters, and LymphGen unclassified DLBCLs (∼40% of all DLBCLs) were broadly distributed among all four IQs **(Figure 3A)**.

**Figure 3.**
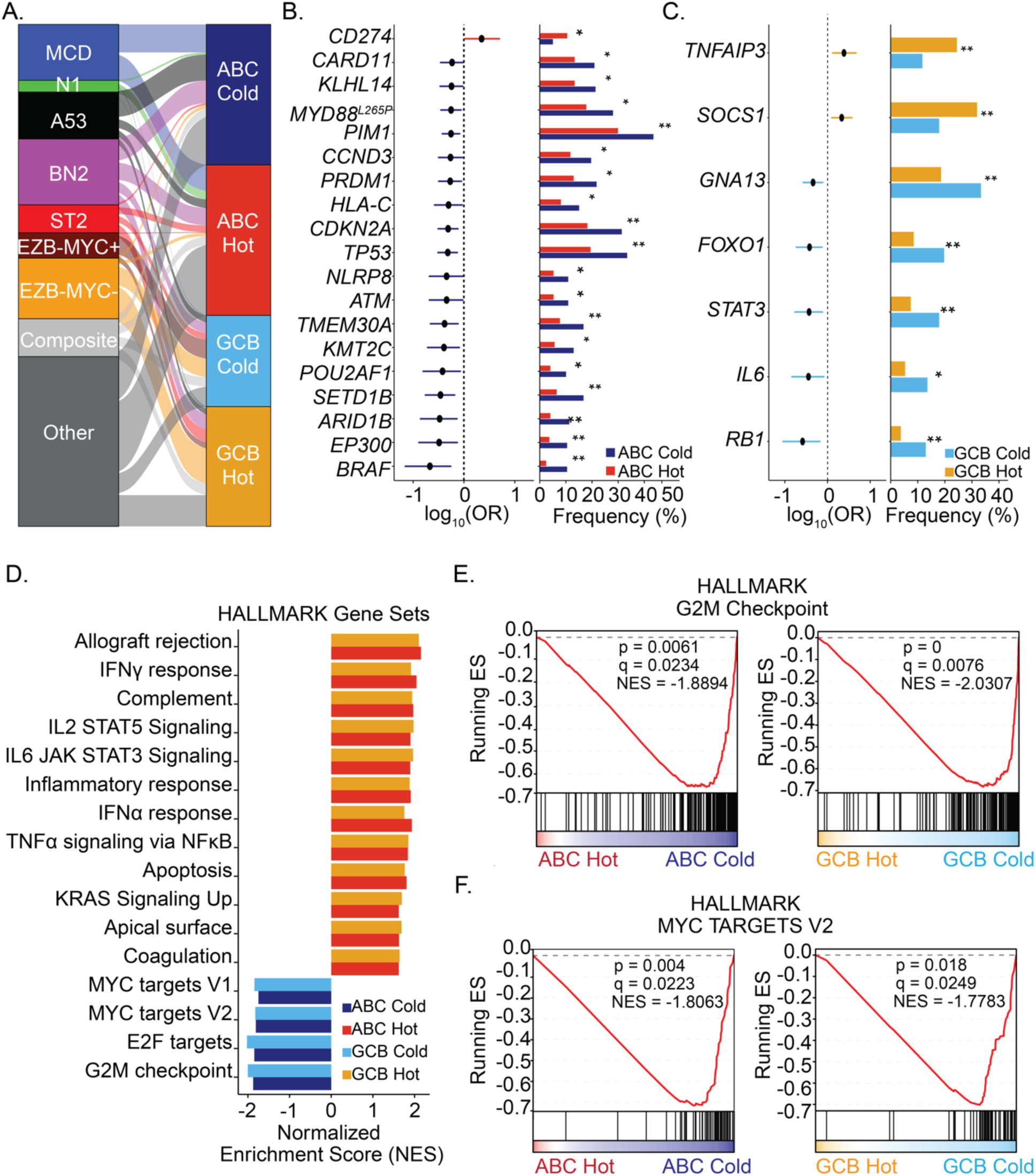
Genomic features associated with DLBCL-IQs. **A.** Alluvial plot showing overlap between LymphGen subtypes and DLBCL-IQs. **B.** Forest plot showing genetic alterations recurrently associated with ABC Hot or ABC Cold DLBCLs. **C.** Forest plot showing genetic alterations recurrently associated with GCB Hot or GCB Cold DLBCLs. **D.** Bar plot of gene set enrichment analysis (GSEA) showing gene sets significantly enriched in each immune-related cluster. **E.** GSEA plots showing upregulation of G2M target gene sets in ABC cold and GCB cold DLBCLs. **F.** GSEA plots showing upregulation of MYC checkpoint related genes in ABC cold and GCB cold DLBCLs. OR – odds ratio; ES – enrichment score; NES – normalized enrichment score. Fisher’s exact test with BH-adjusted p values displayed. (* adj. p < 0.1, ** adj. p < 0.05, *** adj. p < 0.01).

To examine whether individual mutations or CNAs were associated with particular DLBCL-IQs, we compared the genetic landscapes of DLBCLs assigned to each of the four DLBCL-IQs (n = 842). The analysis was restricted to 190 essential driver genes to identify those most relevant to DLBCL biology **(Supplementary Table 2)** (38,40). Comparative analysis of hot DLBCLs versus cold DLBCLs (as defined by PC1) independent of COO identified several genetic lesions associated with cold immune environments, including *FOXO1*, *MYD88,* and *TMEM30A*. Fewer genetic alterations were significantly enriched in hot DLBCLs, but included alterations in *SOCS1*, *TNFRSF14,* and *CD274* **(Supplementary Figure 6A)**. As ABC and GCB DLBCLs arise in the context of distinct genetic alterations and oncogenic pathways, we next compared oncogenic alterations specifically enriched in ABC hot versus ABC cold DLBCLs, and GCB hot versus GCB cold DLBCLs. Alterations in genes such as *MYD88, KLHL14, CARD11,* and *TMEM30A,* known to be potent drivers of chronic BCR signaling, were strongly enriched in ABC cold DLBCLs (41–43) **(Figure 3B)**. Alterations in *CD274* (*PD-L1*) were significantly enriched among ABC hot DLBCLs **(Figure 3B)**, concordant with our previous work (20). Mutations in *RB1*, *FOXO1*, and *GNA13* were enriched in the GCB cold cluster, while those in *TNFAIP3* and *SOCS1* were significantly associated with the GCB hot cluster **(Figure 3C)**.

To gain insight into the impact of oncogenic signaling pathways on DLBCL immune environments, mutations and CNAs were grouped into functional pathways. Notably, genetic alterations in the BCR-dependent NF-κB signaling pathway were significantly enriched in ABC cold DLBCLs compared to ABC hot DLBCLs **(Supplementary Figure 6B)** (43,44). ABC and GCB cold DLBCLs were both significantly enriched for alterations in genes involved in p53 signaling and cell cycle progression compared to ABC and GCB hot DLBCLs, respectively (Supplementary Figure 6C) (**45**).

Finally, gene set enrichment analysis (GSEA) showed upregulation of immune-related gene sets, including IFNγ targets, IFNα response genes, and IL2-STAT5 signaling pathway genes in ABC hot and GCB hot clusters (**Figure 3D**). In contrast, G2M target gene sets were enriched in ABC and GCB cold DLBCL clusters, suggesting dysregulated lymphoma cell cycling might play a role in orchestrating cold DLBCL immune environments **(Figure 3D and E)**. GSEA also demonstrated that ABC and GCB cold DLBCLs were significantly enriched for expression of MYC target genes compared to ABC and GCB hot DLBCLs (**Figure 3D and F)**. Taken together, these data show that several genetic lesions and oncogenic pathways are recurrently associated with specific GSVA-defined DLBCL-IQs, suggesting that lymphoma cell-intrinsic alterations may impact the immune environment in DLBCL.

### MYC activation is strongly associated with cold DLBCL immune environments

Because GSEA showed striking upregulation of MYC targets in cold DLBCLs, the role of MYC in orchestrating a cold immune environment was further interrogated. We devised a MYC GSVA score using genes from a previously published target gene signature (MycUp-4) to assign DLBCLs into “MYC-High,” “MYC-Low”, or “MYC-Int” groups (46). IHC for MYC expression was performed on DLBCLs in the BCC cohort with paired transcriptomic data (n = 266). DLBCLs with high MYC protein expression – defined as ≥ 40% of lymphoma cells expressing MYC by IHC staining – were consistently classified as MYC-high transcriptionally (MYC-High). Conversely, DLBCLs identified as MYC-Low transcriptionally were characterized by low MYC protein expression. Thus, expression of the transcriptional MYC signature accurately identifies DLBCLs with high versus low MYC protein expression **(Supplementary Figure 7A)**.

ABC cold and GCB cold DLBCLs showed significantly higher expression of the MYC activation signature compared to ABC hot or GCB hot DLBCLs **(Supplementary Figure 7B)** and MYC-High DLBCLs were equally distributed between ABC cold (40%) and GCB cold (40%) clusters **(Supplementary Figure 7C)**. Compared to their MYC-Low counterparts, MYC-High DLBCLs in both ABC and GCB clusters contained significantly lower inferred proportions of immune cell subsets when analyzed by CIBERSORTx **(Supplementary Figure 7D-F)**, suggesting a negative correlation between MYC activation and an inflamed environment (47). Supporting this notion, DLBCLs in the MYC-High group were characterized by lower CD8^+^ T cell to DLBCL cell ratios compared to MYC-Low DLBCLs when assessed by mIF **(Supplementary Figure 7G)**. CD8^+^ T cell to DLBCL cell ratios were also negatively correlated with a sample wise MYC GSVA score **(Supplementary Figure 7H)**. Similar findings were also observed for CD4^+^ T cells **(Supplementary Figure 7I and J)**. Taken together, these data reveal a strong association between MYC activity and cold immune environments in DLBCL.

### *SOCS1* mutations are enriched among GCB hot DLBCLs and enhance B-cell sensitivity to IFNγ

We were particularly intrigued by the strong association between GCB hot DLBCLs and putative LoF *SOCS1* mutations as SOCS1 is also frequently mutated in classical Hodgkin lymphoma (cHL) (48–50) and primary mediastinal B cell lymphoma (PMBL) (48,51–53) – two lymphoma subtypes associated with inflamed immune environments and remarkable sensitivity to CBT (16,17,54–57). SOCS1 is a tumor suppressor and potent negative regulator of JAK/STAT signaling, and LoF mutations in *SOCS1* have been implicated in enhancing oncogenic JAK/STAT signaling in lymphoma downstream of cytokines such as IL-4 and IL-13 (58–62). However, as critical T cell effector cytokines such as IFNγ also signal through the JAK/STAT pathway, we hypothesized that LoF *SOCS1* mutations may render DLBCLs more sensitive to the effects of IFNγ, which could stimulate the generation of “inflamed” environments (63–65).

*SOCS1* LoF alterations were identified in 16.2% of DLBCLs in our combined cohort and were significantly associated with a GCB COO **(Figure 4A**). As expected, the incidence of *SOCS1* mutations was significantly higher in the GCB hot cluster (30.3%) compared to other IQs **(Figure 4A and B)**. *SOCS1* mutations were often missense or nonsense (82.3%) and occurred with similar frequencies across the *SOCS1* gene **(Figure 4B),** consistent with the role of SOCS1 as a tumor suppressor. Immune cell deconvolution revealed an increased inferred fraction of several immune cell subsets, including CD8^+^ T cells and CD4^+^ T cells, in *SOCS1* mutant compared to *SOCS1* wildtype GCB DLBCLs suggesting that GCB DLBCLs with *SOCS1* mutations are more inflamed than their *SOCS1* wildtype counterparts **(Figure 4C)**. Moreover, among GCB DLBCLs, GSEA demonstrated increased expression of IFNγ and IFNα signaling pathway associated genes in *SOCS1* mutant DLBCLs compared to *SOCS1* WT DLBCLs **(Figure 4D)**.

**Figure 4.**
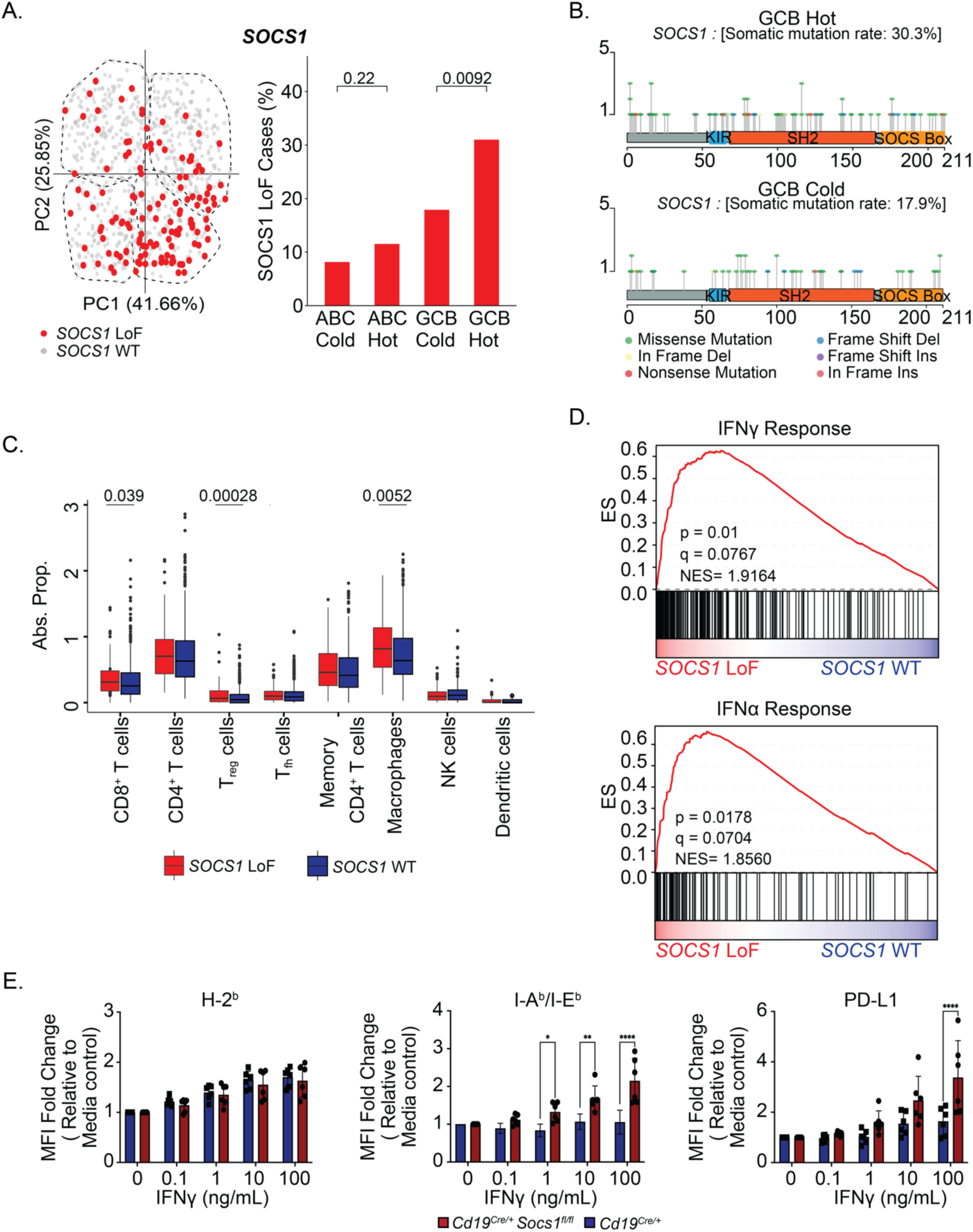
*SOCS1* mutations are enriched among GCB Hot DLBCLs and enhance B-cell sensitivity to IFNγ signaling. **A.** PCA plot (left) and bar plot (right) showing the frequency of *SOCS1* loss of function (LoF) alterations in each immune-related cluster. **B.** Lollipop plots showing frequency and distribution of mutations in *SOCS1* in GCB Hot (top) and GCB cold (bottom) DLBCLs. **C.** Immune cell deconvolution (CIBERSORTx) showing absolute inferred proportions of indicated immune cell subsets in *SOCS1* LoF GCB DLBCLs compared to *SOCS1* wildtype (WT) GCB DLBCLs. **D.** GSEA plots showing upregulation of IFNγ response genes and IFNα response genes in *SOCS1* LoF GCB DLBCLs compared to *SOCS1* WT GCB DLBCLs. Fisher’s exact test with BH-adjusted p values for categorical variables. Kruskal-Wallis test followed by post-hoc Dunn’s test with adjusted p values for continuous variables. **E.** Fold change in mean fluorescence intensity (MFI) of H-2^b^, I-A/I-E^b^, and PD-L1 on CD19^+^ splenocytes. CD19^+^ splenocytes from *Cd19^Cre/+^* (n = 6) or *Cd19^Cre/+^ Socs1^fl/fl^* mice (n = 6) were cultured with media or the indicated concentrations of IFNγ for 48 hours and expression levels of H-2^b^, I-A/I-E^b^, and PD- L1 were measured. Mice were pooled from 3 independent biological replicates. Two-way ANOVA with Bonferroni correction, adj. p values displayed (* adj. p < 0.05, ** adj. p < 0.01, *** adj. p < 0.001, **** adj. p < 0.0001).

To directly test the hypothesis that loss of Socs1 function would render B cells more sensitive to IFNγ, splenocytes from *Socs1*-deficient (*Cd19*^Cre/+^*Socs1^fl/fl^)* or *Socs1*-sufficient mice (*Cd19*^Cre/+^) were subjected to IFNγ stimulation. Representative gating of CD19^+^ B cells and CD3^+^ T cells is shown in **Supplementary Figure 8A**. As hypothesized, IFNγ response genes, such as MHC-II, and PD-L1, were significantly more inducible by IFNγ in a dose-dependent manner in splenic CD19^+^ B cells from mice lacking *Socs1* compared to wildtype controls **(Figure 4E, Supplementary Figure 8B)**. This effect was specific to CD19^+^ cells in which *Socs1* had been deleted, as MHC-II and PD-L1 were similarly inducible in CD3^+^ T cells isolated from spleens of conditional *Socs1*-sufficient and *Socs1*-deficient mice **(Supplementary Figure 8C)**. These data demonstrate that DLBCLs with *SOCS1* mutations represent a novel subset of inflamed lymphomas that may be particularly vulnerable to T cell-based immunotherapies due to enhanced IFNγ sensitivity.

### DLBCL-IQs are associated with distinct survival outcomes to BsAb therapy but not CAR T cell treatment

The presence of activated T cells in the tumor environment has been associated with response to CBT in solid cancers (8–11,66,67). However, the extent to which the local immune environment impacts the effectiveness of immunotherapies, such as BsAbs and CAR T cells, has not been well characterized in DLBCL. Therefore, we sought to define the degree to which our DLBCL-IQs correlated with clinical benefit following treatment with mosunetuzumab, a CD20 x CD3 BsAb and axicabtagene ciloleucel (axi-cel) CD19 CAR T cell therapy.

First, using the immune- and COO-related gene sets detailed earlier, GSVA was performed on the transcriptomes of DLBCLs in our merged dataset (NCI/UCMC/BCC), along with pre-treatment r/r DLBCL biopsies (n = 74) from patients enrolled in a phase I/II dose-escalation trial of mosunetuzumab (5). Consistent with observations in our training and validation datasets, PCA again revealed that PC1 reflected an immune-related axis and PC2 reflected a COO-related axis, and segregated DLBCLs into four quadrants **(Figure 5A and B).** Next, we addressed the extent to which GSVA-defined DLBCL-IQs were associated with differential outcomes to mosunetuzumab treatment. Independent of COO, r/r DLBCLs characterized as “hot” (n = 32) demonstrated significantly improved PFS with mosunetuzumab treatment (HR = 0.58 (0.34-0.97), p = 0.038) compared to those classified as “cold” (n = 42) **(Figure 5C)**. Outcomes to mosunetuzumab treatment were next compared between ABC hot and ABC cold, as well as GCB hot and GCB cold DLBCLs. Interestingly, patients with GCB hot DLBCLs demonstrated significantly improved PFS compared to those with GCB cold DLBCLs (HR = 0.35 (0.14-0.85), p = 0.02) **(Figure 5D).** Patients with GCB cold DLBCLs demonstrated extremely poor outcomes after treatment with mosunetuzumab, with a median PFS of only 2.3 months. Conversely, PFS following mosunetuzumab treatment was similar among patients with ABC hot and ABC cold DLBCLs, indicating that the pre-treatment immune environment can significantly influence response to mosunetuzumab immunotherapy in patients with r/r GCB DLBCL (**Supplementary Figure 9A)**.

**Figure 5.**
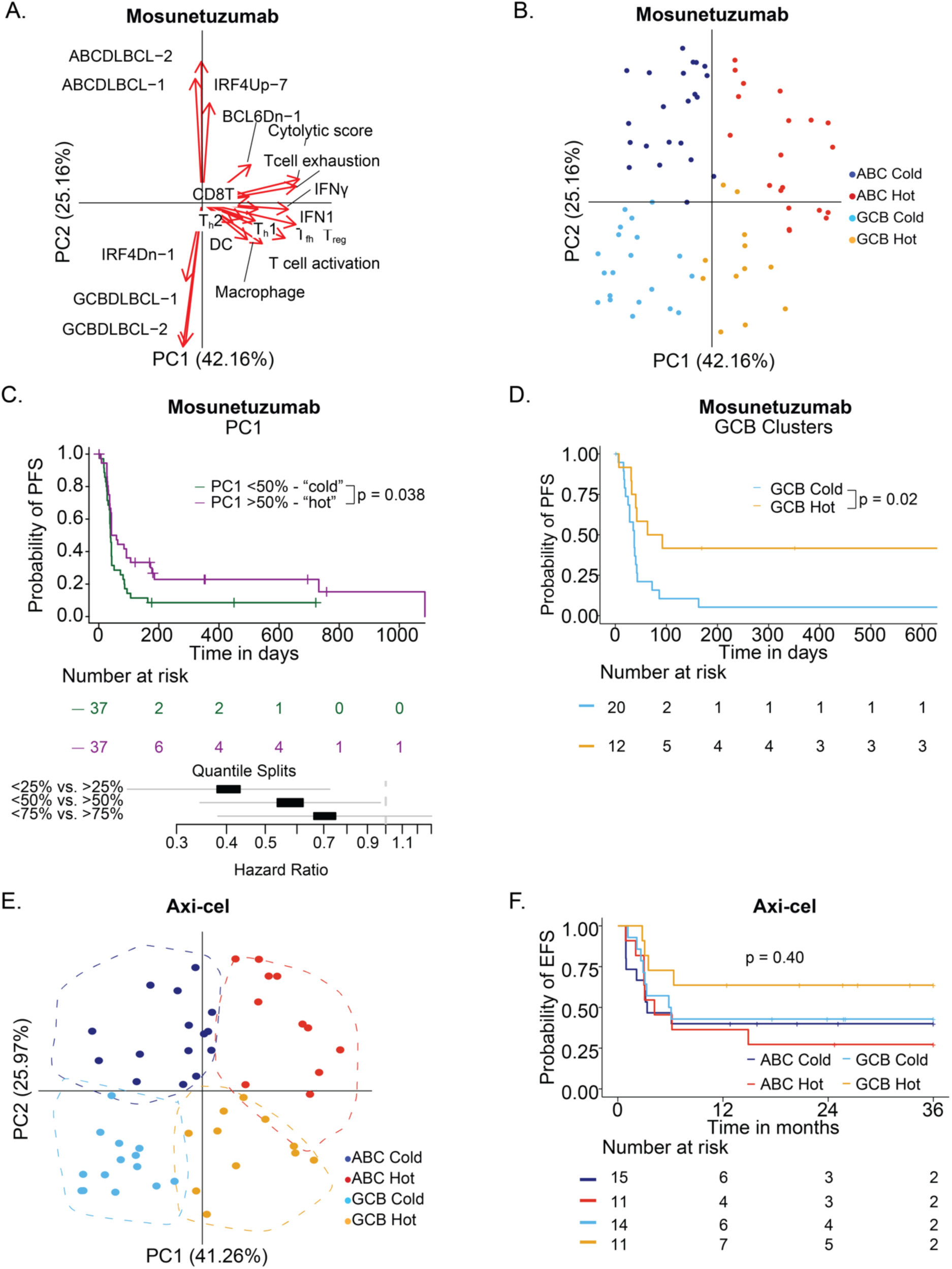
DLBCL-IQs are associated with distinct survival outcomes to BsAb therapy but not CAR T cell treatment. **A.** PCA biplot showing the contribution of immune-related gene sets and COO-related gene sets to PC1 and PC2, respectively, for pre-treatment r/r DLBCL biopsies (n = 74) from patients treated with mosunetuzumab. **B.** PCA plot showing sample-wise GSVA enrichment scores for DLBCL biopsies from patients treated with mosunetuzumab. **C.** PFS for patients assigned to “hot” or “cold” DLBCL immune clusters and treated with mosunetuzumab (top) and forest plot showing hazard ratio of the association between PC1 score and PFS (bottom). **D.** PFS for patients assigned to GCB Hot and GCB Cold DLBCL IQs and treated with mosunetuzumab. Log-rank test with p value displayed. **E.** PCA plot showing sample-wise GSVA enrichment scores for DLBCL biopsies from patients treated with axicabtagene ciloleucel (axi- cel). **F.** Event free survival (EFS) for patients assigned to each DLBCL-IQ and treated with axi- cel. Log-rank test with p value displayed.

As DLBCL-IQs were associated with differential clinical benefit with mosunetuzumab, we hypothesized this might also be the case for other T cell-based immunotherapies, such as CAR T cell therapy. Therefore, we performed GSVA on pre-treatment biopsies of DLBCL patients treated with axi-cel (n = 51) combined with our merged dataset (NCI/BCC/UCMC) (68). Similar to our previous results, DLBCLs were distributed along an immune-related axis (PC1) and COO-related axis (PC2) into four equally sized IQs **(Supplementary Figure 9B and Figure 5E)**. However, in contrast to patients treated with mosunetuzumab, patients with “hot” DLBCLs (n = 33, PC1 > 0) had similar event free survival (EFS) and OS as those with “cold” DLBCLs (n = 40, PC1 < 0) following CAR T cell treatment **(Supplementary Figure 9C and D)**. Moreover, specific DLBCL- IQs were not significantly associated with improved EFS or OS with CAR T cell therapy, although DLBCLs classified as GCB hot trended towards improved EFS **(Figure 5F and Supplementary Figure 9E)**. Taken together, these data suggest that the DLBCL immune environment is correlated with benefit to BsAbs but not CAR T cell therapy, indicating that the mechanism and extent to which these immunotherapies rely on the endogenous immune environment may differ.

## Discussion

Previous transcriptional analyses have demonstrated significant heterogeneity in the immune landscapes of DLBCL (12,15,68). Yet, the precise mechanisms governing these distinct immune environments and their potential association with the response to immunotherapy in DLBCL remain largely unexplored. Here, we curated immune- and COO-related gene sets and performed GSVA to analyze bulk transcriptomes of 862 diagnostic DLBCL samples across three large datasets, which effectively segregated DLBCLs into four IQs: GCB cold, GCB hot, ABC cold, and ABC hot. Importantly, we identified strong associations between recurrent genetic alterations and each GSVA-defined IQ, which suggests that tumor cell-intrinsic alterations have a significant impact in regulating the immune landscape of DLBCL. Finally, we uncovered a striking relationship between the pre-existing immune landscape and response to CD20 x CD3 BsAb treatment, but not CAR T cell therapy, in patients with r/r DLBCL.

Interestingly, our data indicate that classifying DLBCLs along an immune axis (PC1) score can accurately identify “inflamed” DLBCLs that exhibit improved PFS to treatment with the CD20 BsAb, mosunetuzumab. Surprisingly, we identified that COO significantly influences this association, as only GCB hot DLBCLs exhibited improved PFS following mosunetuzumab treatment. This finding suggests that for r/r GCB DLBCLs, the pre-existing immune environment is clearly an important factor associated with benefit from BsAb immunotherapy. In contrast, the PC1 axis was not associated with improved survival following treatment with axi-cel. These findings align with a recent analysis of pre-treatment DLBCL biopsies from patients enrolled in the ZUMA-7 study, where the expression of T cell-associated transcripts was not associated with a significant EFS advantage after second-line axi-cel treatment (69). However, when considering COO (PC2) in addition to PC1, we find that patients with GCB hot DLBCLs treated with axi-cel also demonstrated a trend toward improved EFS compared to DLBCLs assigned to other DLBCL- IQs similar to mosunetuzumab. Together, these data suggest that integrating COO and immune signatures, as with DLBCL-IQ, may provide a more nuanced understanding of the factors that may govern response or resistance to BsAb and CAR T cell therapy.

A central hypothesis underlying our analyses was that lymphoma cell-intrinsic alterations significantly influence the composition of the DLBCL immune environment. Supporting this notion, we found that each DLBCL-IQ was significantly enriched for distinct genetic lesions and oncogenic pathways. For example, GCB hot DLBCLs were significantly enriched for *SOCS1* LoF mutations, which are commonly observed in other inflamed lymphomas that are highly susceptible to CBT, such as cHL and PMBL. Because *SOCS1* is a potent negative regulator of JAK/STAT signaling, we hypothesized that LoF *SOCS1* mutations would sensitize B cells to IFNγ – a JAK/STAT-dependent effector cytokine critical for effective anti-tumor immune responses (8,9,61–63,70). Indeed, genetic alterations that blunt IFNγ sensing in tumors, such as LoF *JAK1/2* alterations, lead to resistance to T cell-based immunotherapies (32,67,71–74). In contrast, loss of negative regulators of IFNγ signaling, for example *PTPN2, PBRM1,* and *ARID2*, sensitize tumor cells to IFNγ and T cell mediated cytolysis, are associated with inflamed tumor environments, and have been linked with CBT responsiveness in solid tumors (34,35,75,76). Although the extent to which LoF mutations in *SOCS1* directly mediate sensitivity to CBT in cHL and PMBL remains to be elucidated, we find that genetic ablation of *Socs1* sensitizes murine B cells to IFNγ. Therefore, these data suggest that *SOCS1* mutations identify a novel subset of “inflamed” DLBCLs that may be inherently vulnerable to T cell-based immunotherapies that rely on this key effector cytokine.

Among ABC DLBCLs, mutations culminating in enhanced BCR activation, such as *MYD88^L265P^, CARD11, KLHL14*, and *TMEM30A* were significantly associated with cold immune environments (26,44,77,78). The potential mechanism(s) by which oncogenic BCR signaling in lymphoma cells leads to decreased T cell infiltration is unknown. However, we observed that ABC cold DLBCLs exhibited higher MYC activity than GCB cold DLBCLs – a surprising finding since MYC activity in GCB DLBCL is often driven by *MYC* translocations to the *IGH* locus, leading to constitutive MYC expression (79–81). Therefore, we hypothesize that ABC DLBCL-associated oncogenic pathways, namely BCR-dependent NF-κB signaling, may drive particularly potent MYC activity and the acquisition of cold immune environments. This finding would have significant translational relevance, as therapies that indirectly modulate MYC activity by acting on BCR signaling may shift the balance toward a “hot” environment and sensitize DLBCLs to immunotherapies.

While these results represent an important advance in our understanding of the DLBCL immune environment, there are some limitations to our study. Firstly, GSVA assigns relative enrichment scores, which might exhibit variability across datasets, thereby complicating comparisons between different datasets. To address this limitation, we combined the mosunetuzumab and Stanford CAR T cohorts with our merged dataset of DLBCLs (NCI/BCC/UCMC), which may facilitate more direct comparisons between patients treated with mosunetuzumab and axi-cel. Second, our analysis utilized bulk transcriptomic data from diagnostic DLBCL samples, and it is unclear how prior therapies impact the immune landscape in this disease. However, when we applied our GSVA-based clustering to the transcriptomes of two independent cohorts of heavily pre-treated patients with r/r DLBCL, we identified a similar distribution of immune-related scores, which preliminarily suggests a relatively limited effect of cytotoxic therapy on the composition of DLBCL immune environments. Finally, the association between particular DLBCL-IQs and clinical benefit from mosunetuzumab and CAR T cell therapy was based on relatively small patient cohorts and the analyses were retrospective in nature. Despite these limitations, our study provides valuable insights into the DLBCL immune environment and its relevance to therapeutic responses, paving the way for further investigation and refinement of treatment strategies in this complex disease.

In conclusion, our results demonstrate that genomic alterations in lymphoma cells can impact the DLBCL immune environment. Furthermore, we show using DLBCL-IQ the differential impact of the pre-treatment DLBCL immune environment on clinical benefit after BsAb versus CAR T cell therapies. This latter observation is particularly important in suggesting that the effectiveness of BsAbs might require a more “inflamed” microenvironment, whereas CAR T cells may be efficacious even in “cold” DLBCLs. Finally, our data suggest that profiling the DLBCL immune environment could be useful in delivering more personalized immunotherapy to r/r DLBCL patients.

## Methods

### Data sets

#### NCI cohort

RNA-sequencing counts for 481 DLBCL biopsies were downloaded from dbGaP (phs001444) (18). All patients had variants calls from whole exome sequencing and copy number arrays.

#### UCMC cohort

RNA-sequencing was performed on 96 DLBCL biopsies (84 treatment-naïve, 12- relapsed/refractory, 2 unknown). Paired whole exome sequencing was successful for 76 cases.

#### BCC cohort

RNA-sequencing counts for 285 treatment-naive DLBCL biopsies was provided by BCC (27,28). Of these, 283 patients had available variant calls from targeted sequencing and copy number arrays.

#### Stanford cohort

RNA-sequencing data quantified as transcripts per million (TPM) for 88 DLBCL biopsies were downloaded from dbGaP (phs003145.v1.p1) (68).

Data were quantified as log2(TPM+1) for analysis.

### Molecular and genetic subtype classifications

#### Cell of origin

COO classifications were compiled from previously described gene expression-based classifiers. For the NCI cohort, all DLBCLs had COO calls from a DGE-based classifier (25). For the BCC cohort, 283 DLBCLs had COO calls from Lymph3Cx (27). For the UCMC cohort, all samples had COO calls from an RNAseq-based classifier from BostonGene (15).

#### LME classifications

Lymphoma Microenvironment (LME) classifications for DLBCLs in the NCI, BCC, and UCMC cohorts were provided by BostonGene (15).

#### Dark zone signature

DZsig (formerly known as double hit signature) classifications for the BCC and NCI cohorts were obtained from Ennishi et al. (28) and Wright et al. (18), respectively. DZsig classifications for the UCMC cohort were provided by BostonGene from an RNAseq-based classifier (18).

#### *LymphGen* classifications

LymphGen classifications for the BCC and NCI cohorts were obtained from Wright et al. (18).

### Generation and processing of sequencing data

#### Quality control

FastQC and FastQ Screen were used to assess read quality and other species contamination (www.bioinformatics.babraham.ac.uk). SNP (single nucleotide polyphorism)- calling using pileup in FACETS was used to assess sample mix-ups and contamination (82).

#### Mutation-calling and annotation

Whole exome sequencing data for the UCMC cohort was generated as previously described (20). Mutation-calling for whole exome sequencing data for the UCMC cohort was performed similarly to the analysis in Kotlov et al (15). Reads were trimmed using fastp to eliminate low-quality reads and then were aligned to the GRCh38 reference genome using BWA (v0.7.17) (83,84). Duplicate reads were removed using Picard’s MarkDuplicates (v2.6.0) (“Picard Toolkit”, 2019. Broad Institute, GitHub Repository) and recalibrated by BaseRecalibrator and ApplyBQSR from GATK4 (85). For samples with a paired normal reference, short variants were called using Strelka (86). For tumor-only samples, Pisces (v5.2) was used (87). Variants were annotated using Variant Effect Predictor (VEP) (88). For all data sets, variants were restricted to these predicted to be deleterious.

#### Copy number analysis

Copy number variation (CNV) was assessed using Sequenza with modifications from FACETS (82,89). CNV analysis was performed in the space of genes listed in **Supplementary Table 2.** Sequenza input reference was modified to match FACETS, calculated using facets-pileup. For tumor-only samples, calling versus average normal or tumor sample was performed using a modified version of Sequenza. Average normal/tumor pileup files were prepared to average normalized coverage and variant allele fraction (VAF) at each position. Some samples were excluded due to low coverage and/or tumor content. For all data sets, heterozygous deletions and low-level copy gains were excluded.

#### RNA-seq processing and normalization

Paired-end and single-end RNA-seq data for the UCMC cohort was generated as previously described (20). RNA-seq reads were aligned using kallisto v0.42.4 to GENCODE v23 transcripts with default parameters (90). The protein-coding transcripts, immunoglobulin heavy, kappa and lambda light chains, and TCR-related transcripts were retained; noncoding RNA, histone, and mitochondria-related transcripts were removed. Counts were processed using limma (v3.52.4) and edgeR (v3.38.4). Genes with low counts were removed using a threshold of a 0.5 log2CPM average across all samples, as previously described (21). Then, TMM (trimmed mean of M-values) normalization was performed, and counts were quantified as log2CPM. Batch correction was performed using the removeBatchEffect function in limma. For DLBCLs in the mosunetuzumab cohort, robust library sizes were estimated using DESeq2, and raw RNASeq counts were normalized via limma-voom.

### Differential gene expression (DGE) and gene set enrichment analysis (GSEA)

DGE was performed using voom, and differentially expressed genes were generally defined as those with |log2FC| ≥ 1 and adjusted p value < 0.05 (91). GSEA was performed using the Desktop client (v4.2.3) with default settings (92).

### Comparing variants between groups

Variants were restricted to previously described DLBCL driver genes (38,40), listed in **Supplementary Table 2.** Maftools (v2.12.0) was used to compare alterations between groups, where additional filtering was performed to filter to genes altered in at least 10% of samples of the smaller group (93).

### Mutational pathway analysis

Genes in the BCR-dependent NFKB pathway (44,45) and cell cycle pathway-related genes (45) were curated from previously published studies. The proportion of patients with an alteration in at least one gene in the pathway was calculated and compared across clusters. Alterations for a given gene were filtered to reflect their putative role as an oncogene or as a tumor suppressor gene.

### CIBERSORTx

Immune cell deconvolution was performed using CIBERSORTx using the LM22 signature matrix(47,94). The software was run on bulk-mode and absolute mode. For the input mixture file, RNA-seq counts were first batch corrected using ComBat-Seq followed by quantification to transcripts per million (TPM) (95).

Cell types were condensed as such:

**T.cells** = “T cells CD8” + “T cells follicular helper (T_fh_)” + “T cells regulatory (Tregs)” + “T cells CD4 naïve” + “T cells CD4 memory resting” + “T cells CD4 memory activated”+ “T cells gamma delta”

**NK.cells** = “NK cells activated” + “NK cells resting”

**Memory.CD4.T.cells** = “T cells CD4 memory resting” + “T cells CD4 memory activated” **Conv.CD4.T.cells** = “Memory.CD4.T.cells” + “T cells CD4 naïve” + “T cells follicular helper” **CD4.T.cells** = “Memory.CD4.T.cells” + “T cells CD4 naïve” + “T cells follicular helper” + “T cells regulatory (T_regs_)”

**Dendritic.cells** = “Dendritic cells activated” + “Dendritic cells resting”

**B.cells** = “B cells naïve” + “B cells memory” + “Plasma cells”

**Macrophages** = “Macrophages M0” + “Macrophages M1” + “Macrophages M2”

EcoTyper (https://github.com/digitalcytometry/ecotype) was used to recover previously defined Lymphoma Ecotypes; input data were quantified as TPM (96).

### Gene set compilation

Gene sets were manually curated following an extensive literature search and focused on genes related to 1) immune cell infiltration and activation, 2) cell-of-origin and 3) key transcription factors related to cell-of-origin. Gene sets were excluded if they had fewer than 3 or more than 200 genes. Genes included in each gene set are listed in **Supplementary Table 1**.

#### Immune-related gene sets

Twelve immune-related gene sets were included, known to be associated with immune activity in other cancer types. Gene sets were related to activation state of T cells (t.cell.activation (11), t.cell.exhaustion (11), IFNγ_ayers (8)), non-cellular mediators of immune responses (ifn1_rooney (11), cytolytic.score (23)), and presence of immune cell subsets (tfh_charoentong (22), th1_charoentong (22), th2_charoentong (22), treg_rooney (11), cd8t_charoentong (22), macrophage_rooney (11), dc_xcell_total (24)). In general, immune-related gene sets had less than 15% overlap based on pairwise comparisons.

#### Cell of origin gene sets

A total of four COO gene sets - 2 ABC DLBCL (ABCDLBC-1, ABCDLBCL-2) specific gene sets and 2 GCB DLBCL specific gene sets (GCBDLBCL-1, GCBDLBCL-2) - were included. Gene sets are derived from the seminal gene expression classifier (25), found in SignatureDB (SigDB)(46). (https://lymphochip.nih.gov/signaturedb/index.html).

#### Transcription factor gene sets

Three transcription factor related gene sets – related to IRF4 (IRF4Up-7 and IRF4Dn-1) and BCL6 (BCL6Dn-1) targets – were included from SigDB (46).

### Immune signature clustering model

Gene set variation analysis (GSVA) using the 19 curated gene sets was performed on bulk transcriptomes from 2 genomic datasets (NCI, n = 481; UCMC, n = 96) (18,20). The resulting GSVA scores were resolved using principal component analysis (PCA). Unsupervised k-means clustering was utilized to assign samples into unique clusters, where k = 4 clusters was optimally selected using an elbow plot. Subsequent analyses using the third independent BCC dataset as well as the 3-dataset combination were performed in the same structured steps, where equivalent clustering was observed.

### Generation of MYCUp4 expression groups

GSVA scores for the MYCUp-4 gene set (from SigDB (46)) were generated for all DLBCLs in UCMC, NCI, and BCC datasets. DLBCLs were then split into ABC and GCB groups based on their immune-related cluster membership, and a 25-75 percentile split within each COO group was used to assign DLBCLs to “MYC-High” and “MYC-Low” groups. All other DLBCLs were assigned to the “MYC-Int” group.

### Multispectral immunofluorescence

Multi-spectral immunofluorescence (mIF) microscopy was performed on 65 DLBCLs (UCMC) for which paired RNA-seq and GSVA data were available. mIF analysis was performed after staining with fluorescence-labeled antibodies against a T cell panel and myeloid cell panel *(see antibodies for mIF section*). Each slide was scanned using the Vectra Polaris (Akoya Biosciences) imaging platform and the Phenochart software (PerkinElmer). Through Phenochart, at least 5 representative regions of interest per tissue section were acquired as multispectral images at 40x magnification. Watershed segmentation was used to identify nuclei using DAPI staining and cell borders of individual cells in each ROI. The supervised machine-learning algorithm in the inForm software (v. 2.3) was used to classify each cell into specific phenotypes.

1. Slides were divided by panel and by specific features determined during the initial watershed segmentation, including cell border detection, average cell size, autofluorescence, and DAPI strength.
2. Samples were divided into 4 separate groups for each panel. 2 ROIs were chosen per sample to train the machine-learning algorithm in inForm to identify the following phenotypes with the associated markers:

1. **T cells** : CD8^+^ T cell, CD4^+^ T cell, PAX5^+^ DLBCL cell
2. **Myeloid cells** : CD68^+^ Macrophage, CD11c^+^ dendritic cell
3. Once the initial training groups were processed, all ROIs in each group of samples were classified using the matching training cohort.
4. Results of per-slide frequencies of each phenotype were tabulated in R using exported values from inForm using the ‘phenoptr’ package and the representative mIF images were also exported through inForm. Given heterogeneity of tumor and microenvironment composition in each ROI, comparisons were made between total number of each TME cell population/phenotype normalized against the number of DLBCL cells present across all ROIs per slide.

### Antibodies for mIF

| PAX5 | nucleus | 620 | BioCare Medical,CM 207 | BC/24, mouse mAb | AR9 | 1:50 in Da Vinci Green |
| --- | --- | --- | --- | --- | --- | --- |
| CD68 | membrane+cytoplasmic | 570 | BioCare Medical, CM033A | KP1, mouse mAb | AR9 | 1:50 in Da Vinci Green |
| CD11c | membrane | 540 | BioCare Medical, ACI3122A | 5D11, mouse mAB | AR9 | 1:100 in Renoir Red |

| CD4 | membrane | 520 | BioCare Medical, ACI3148A | 4B12, mouse mAb | AR9 | 1:50 in Van Gogh Yellow |
| --- | --- | --- | --- | --- | --- | --- |
| PAX5 | nucleus | 620 | BioCare Medical,CM 207 | BC/24, mouse mAb | AR9 | 1:50 in Da Vinci Green |
| CD8 | membrane | 690 | R&D,<br>NBP232836B | C8/144B,<br>mouse mAb | AR9 | 1:50 |

### *In vitro* IFNγ stimulation of splenocytes

Spleens from *CD19^Cre/wt^* and *CD19^Cre/wt^* ; *Socs1^fl/fl^*mice were obtained from Ari Melnick (Weill Cornell Medicine). Spleens were passed through a 70 µm cell strainer (Corning, 352350) and ground using the plunger of a syringe with 10 mL of PBS to yield single cell suspensions. Splenocytes were then washed twice in 10mL of PBS, followed by lysis of red blood cells in a hypotonic solution and three washes in 10 mL of PBS. 5 x 10^5^ splenocytes from single cell suspensions were incubated with increasing concentrations of recombinant mouse IFNγ (0, 0.1, 1, 10, 100 ng/mL) for 48 hours (Peprotech, 315-05). Cells were then harvested and incubated with a panel of antibodies containing anti-mouse CD19 (Biolegend,115504), anti-mouse CD3d (Biolegend, 100244), anti-mouse H-2 (Biolegend, 125506), anti-mouse I-A/I-E (Biolegend, 107619), and anti-mouse PD-L1 (Biolegend, 124311). Splenocytes were analyzed by flow cytometry for expression of H-2, I-A/I-E, and PD-L1 on CD19^+^ and CD3^+^ cell populations. Mean fluorescence intensity (MFI) fold change for H-2, H-2, I-A/I-E, and PD-L1 was computed by comparing treatment with IFNγ compared to media-only control. Data shown are the average of at least 3 independent biological replicates. Statistical testing was conducted using a 2-way ANOVA with Bonferroni correction for multiple comparisons.

### Statistical methods

All statistical analyses were performed in R (v4.2.1). Statistical significance for categorical variables was assessed with a Fisher’s exact test for two-group comparisons; a multi-way Fisher’s exact test followed by post-hoc individual pairwise testing was used for comparisons of three or more groups. Statistical significance of continuous variables was performed with a Mann-Whitney U test for two-group comparisons and a Kruskal-Wallis test followed by a post-hoc Dunn’s test for comparisons across three or more groups. Survival analysis was performed using the survival package (v3.4.0), and the log-rank test was used to determine significance. A p value threshold of 0.05 was used to assess univariate statistical significance. Adjustment for multiple hypothesis testing was performed using the Benjamini-Hochberg method unless indicated otherwise.

