## Supplementary Figures for "Integrative genomic analysis identifies unique immune environments associated with immunotherapy response in diffuse large B cell lymphoma"

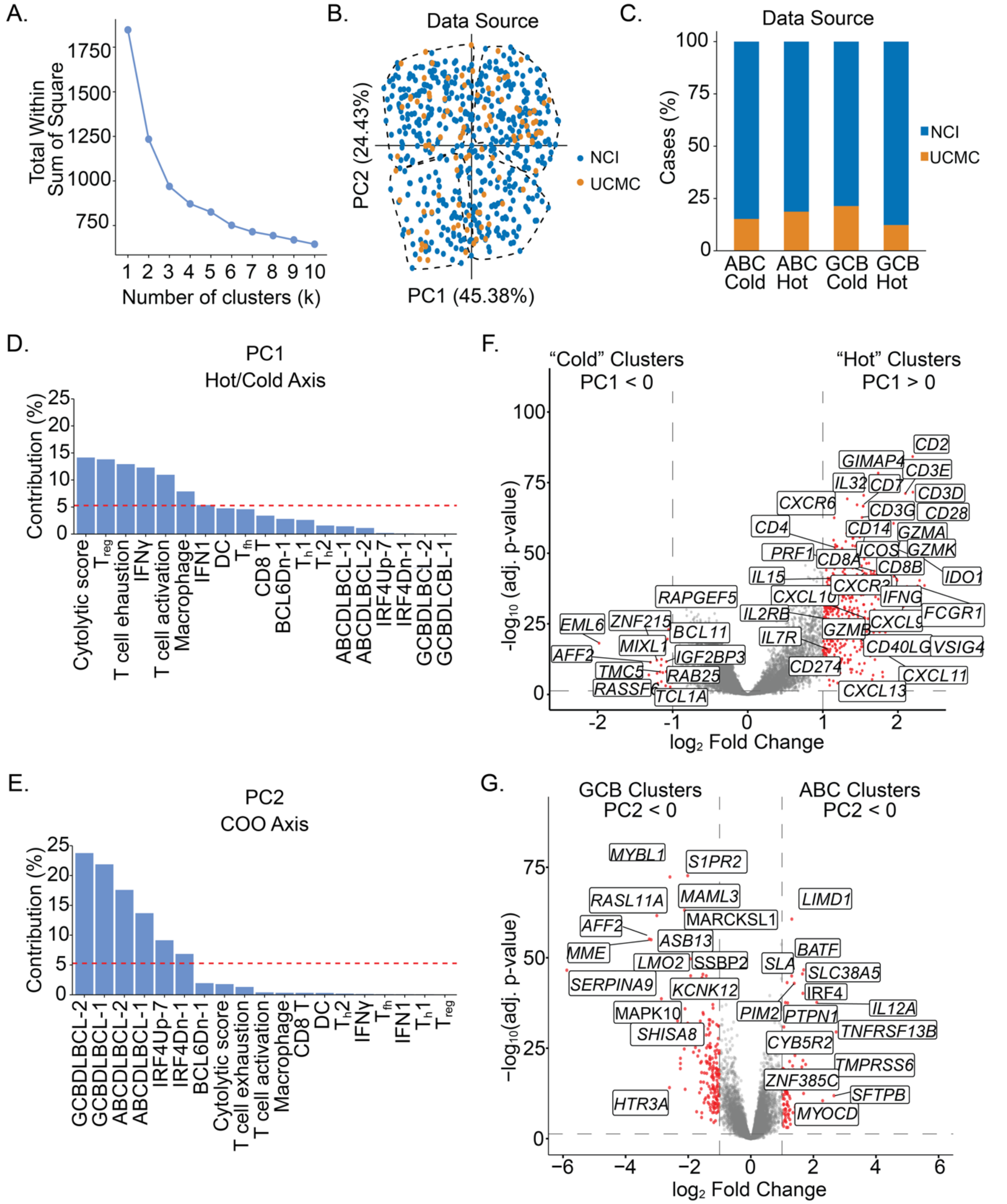


**Supplementary Figure 1. Transcriptomic analysis identifies four unique immune-related clusters.** **Supplementary Figure 1. Transcriptomic analysis identifies four unique immune-**

**related clusters. A.** Elbow plot showing optimal number of clusters in the data (k = 4). **B.** PCA plot showing distribution of DLBCLs from two independent data sources, indicating no batch effect (NCI – blue, UCMC – yellow). **C.** Bar plot quantifying distribution of cases from two independent data sources. **D-E.** Bar plots showing contribution of 19 immune-related and COO-related gene sets to PC1 **(D)** and PC2 **(E)**. **F.** Volcano plot showing differentially expressed genes ($|\log_2 FC| > 1.0$, adj p value $< 0.05$) between putative "hot" DLBCLs (PC1 > 0) and "cold" DLBCLs (PC1 < 0). **G.** Volcano plot showing differentially expressed genes ($|\log_2 FC| > 1.0$, adj p value $<$ 0.05) between putative ABC DLBCLs (PC2 > 0) and GCB DLBCLs (PC2 < 0).

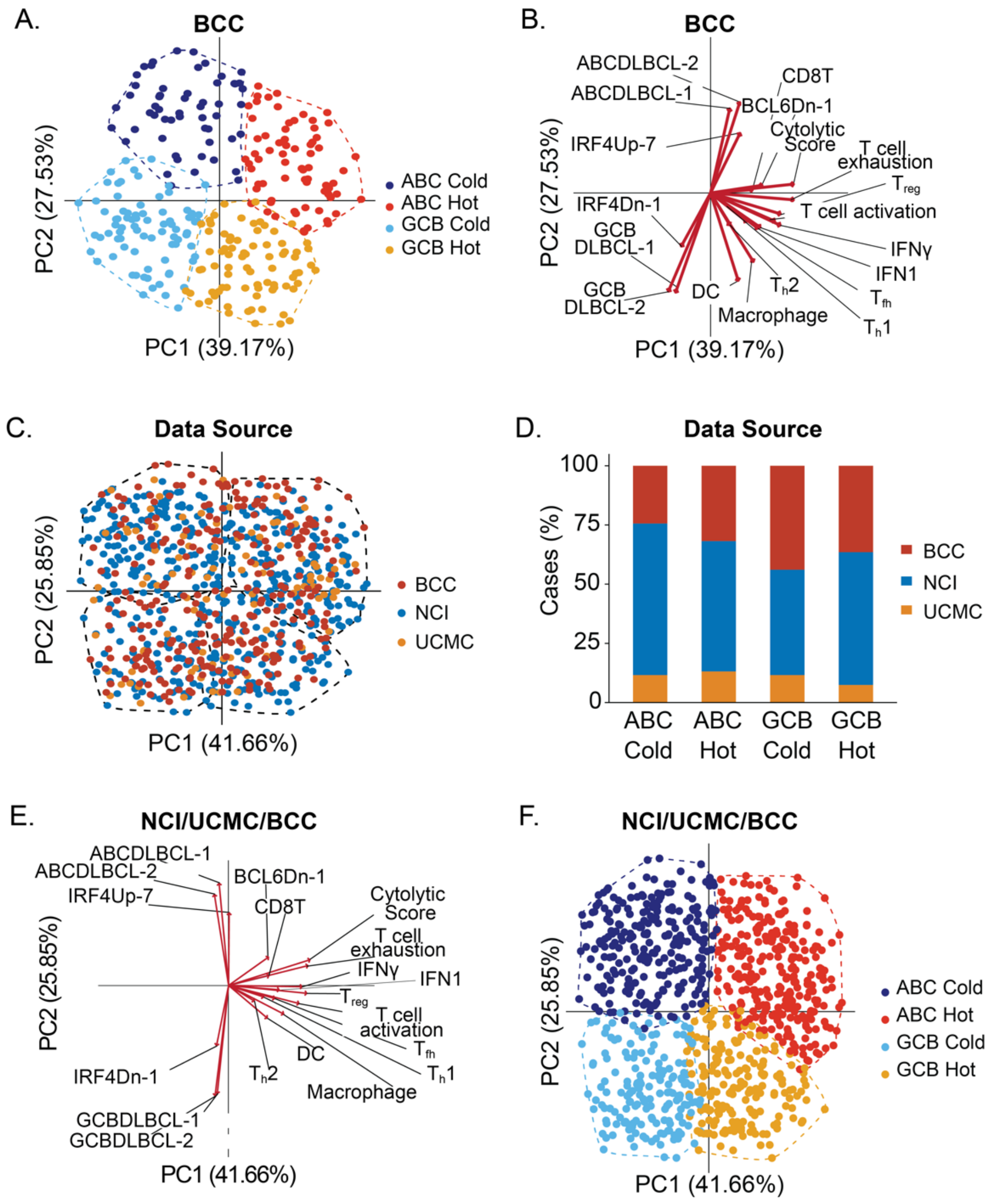


**Supplementary Figure 2. Transcriptomic analysis identifies four unique immune-related clusters. A.** PCA plot showing sample-wise GSVA enrichment scores for DLBCLs in an independent dataset (BCC). **B.** PCA biplot showing contribution of immune-related and COO-related gene sets to PC1 and PC2, respectively, for DLBCLs from the BCC dataset. **C**. PCA plot showing sample-wise GSVA enrichment scores for DLBCLs from all three datasets, colored by data source (NCI – blue, UCMC – yellow, BCC – red). **D.** Bar plot showing distribution of cases from NCI, UCMC, and BCC datasets within each GSVA-based immune cluster. **E**. PCA biplot showing contribution of immune-related and COO-related gene sets to PC1 and PC2 for DLBCLs in the combined NCI/UCMC/BCC dataset. **F**. PCA plot showing sample-wise GSVA enrichment scores for combined NCI/UCMC/BCC dataset.

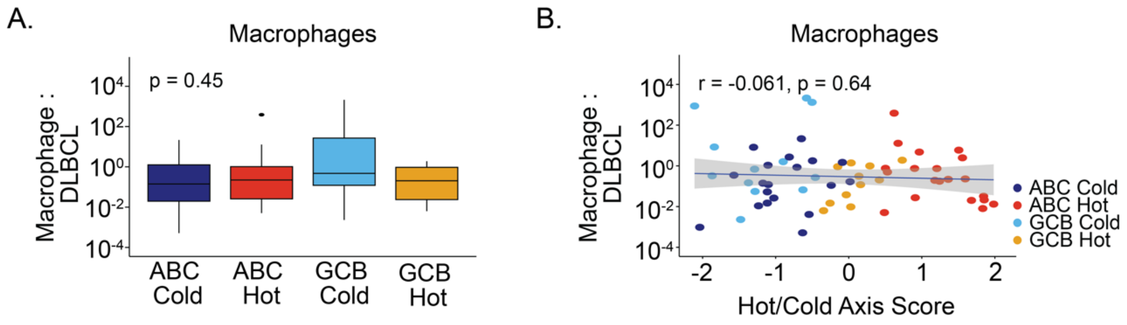


**Supplementary Figure 3. Validation of transcriptionally defined DLBCL-IQs**. **A.** Box plot showing average CD68$^+$ macrophage : DLBCL ratios in each IQ as assessed by mIF (n = 65). **B.** Scatter plot showing correlation of hot/cold axis score (PC1) and CD68$^+$ macrophage : DLBCL ratio (n = 65). Statistical analysis by Kruskal-Wallis test followed by a post-hoc Dunn's test with Benjamini-Hochberg (BH) adjusted p value.

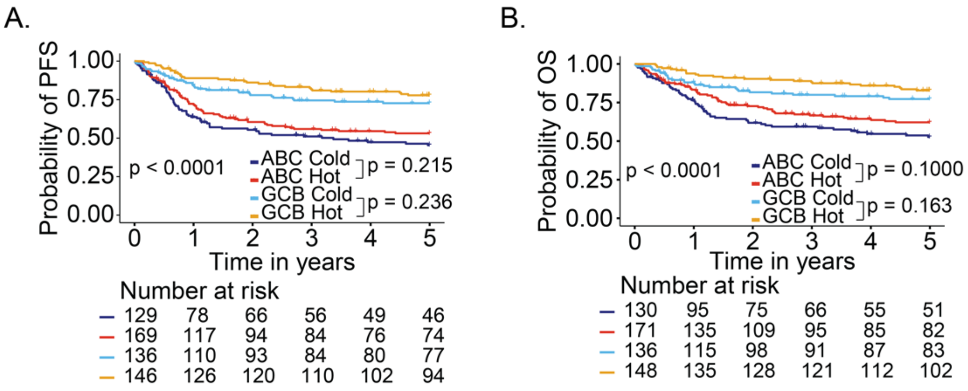


**Supplementary Figure 4. Prognostic significance of DLBCL-IQs. A-B.** Five-year progression free survival (PFS) **(A)** and overall survival (OS) **(B)** for DLBCLs assigned to the indicated IQs. Log-rank test with Benjamini-Hochberg (BH)-adjusted p values.

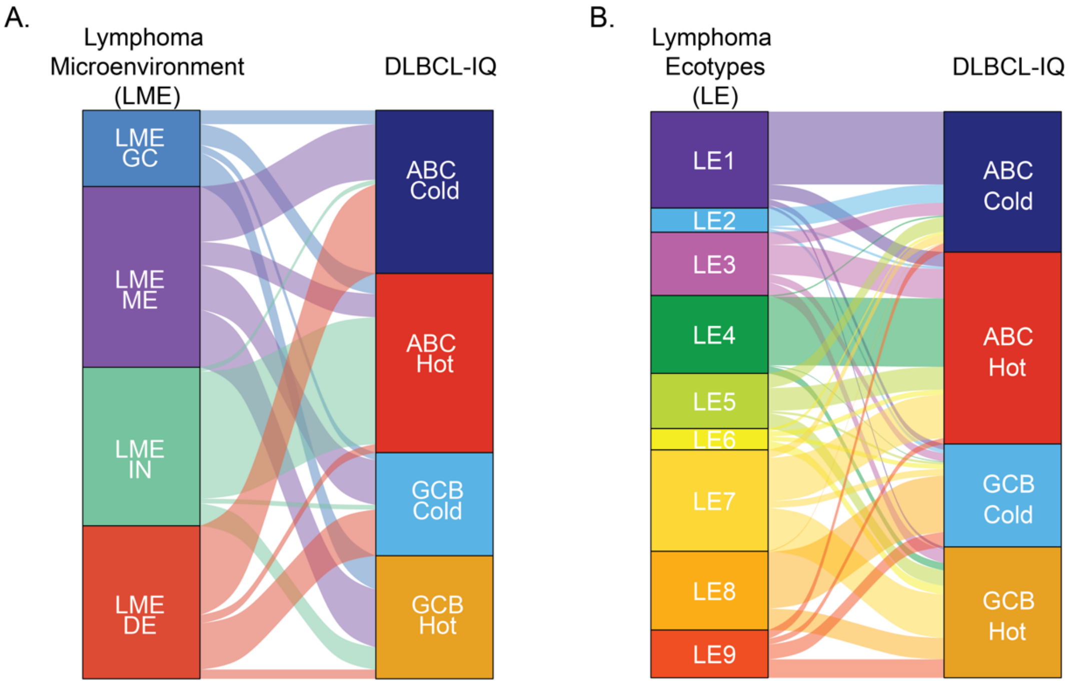


**Supplementary Figure 5. Concordance of DLBCL-IQs with other immune-related DLBCL subtypes.** **A**. Alluvial plot demonstrating overlap between LME clusters and GSVA-defined DLBCL-IQs. **B.** Alluvial plot demonstrating overlap between LE clusters and GSVA-defined DLBCL-IQs. (LME – Lymphoma microenvironment; GC – germinal center; ME – mesenchymal; IN – inflamed; DE – depleted; LE – Lymphoma ecotypes).

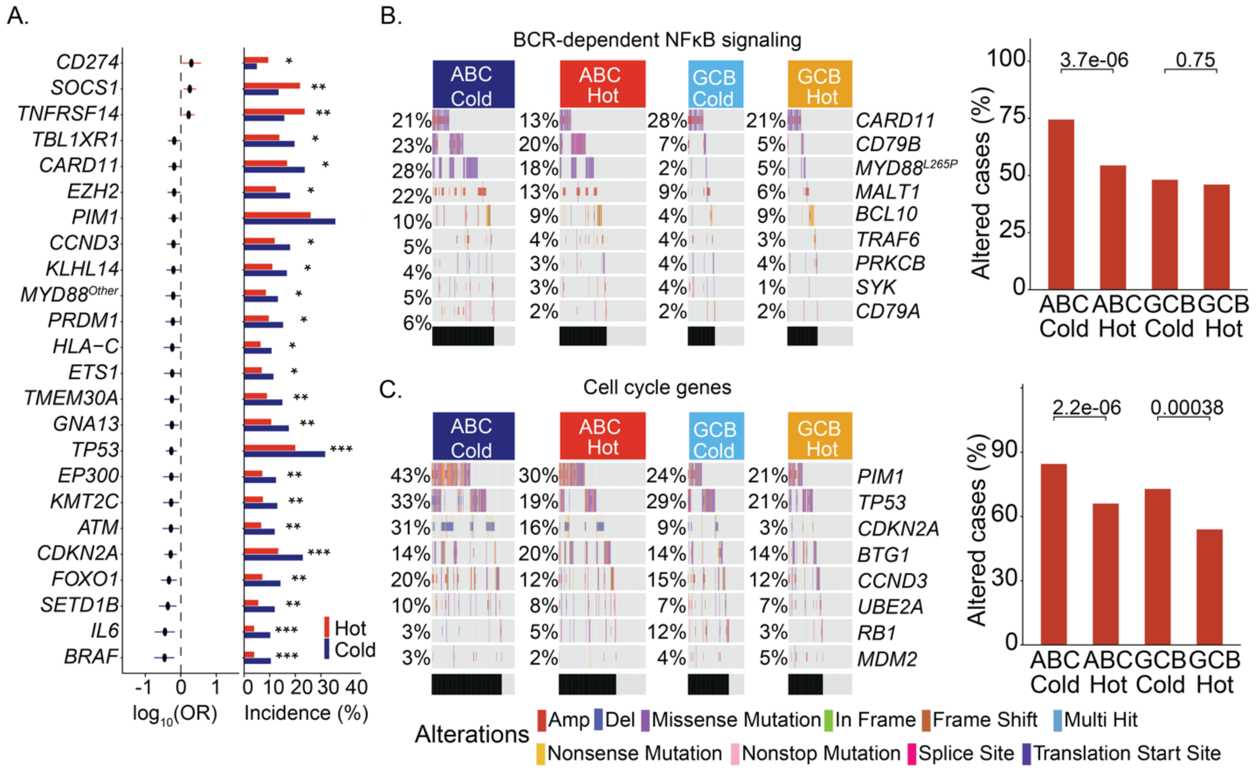


**Supplementary Figure 6. Genomic features associated with DLBCL-IQs. A.** Forest plot of genetic alterations recurrently associated with hot and cold DLBCL-IQs. Fisher's exact test with BH-adjusted p values displayed. (* adj. p < 0.1, ** adj. p < 0.05, *** adj. p < 0.01). **B.** Oncoprint (left) and bar plot (right) showing frequency of alterations in BCR-dependent NFκB pathway genes among the four DLBCL-IQs. **C.** Oncoprint (left) and bar plot (right) showing frequency of alterations in cell cycle genes genes among the four GSVA-defined immune-related clusters. Unadjusted p values displayed.

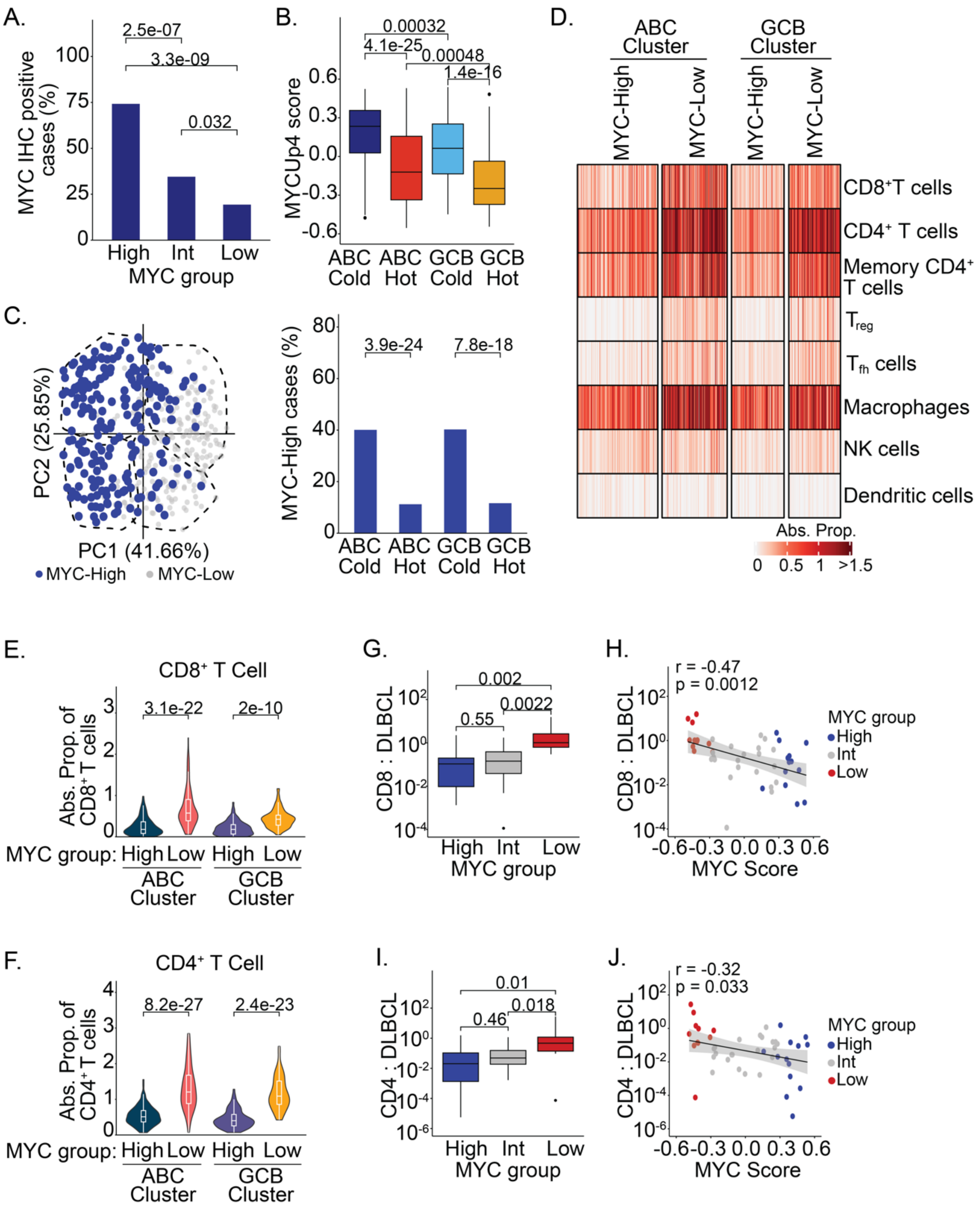


**Supplementary Figure 7. MYC activity is associated with cold DLBCL immune environments. A.** Bar plot showing frequency of MYC IHC+ DLBCLs in transcriptionally defined MYC-High, MYC-Int, MYC-Low groups. **B.** Box plot showing MYC score for DLBCLs in each GSVA-based immune related cluster. **C.** PCA plot (left) and bar plot (right) showing frequency of

MYC-High DLBCLs in immune clusters. **D.** Heatmap showing absolute inferred proportions of immune cell subsets in MYC-High and MYC-Low DLBCLs from ABC and GCB clusters. **E-F.** Violin plots showing absolute inferred proportions of $CD8^+$ **(E)** and $CD4^+$ **(F)** T cells in MYC-High and MYC-Low DLBCLs in ABC and GCB clusters. **G.** Box plot showing $CD8^+$T cell : DLBCL ratio in MYC-High, MYC-Int, MYC-Low groups. **H.** Scatter plot showing correlation of MYCUp4 score and $CD8^+$ T cell : DLBCL cell ratio. **I.** Box plot showing CD4+T cell : DLBCL ratio in MYC-High, MYC-Int, MYC-Low groups. **J.** Scatter plot showing correlation of MYC score and $CD4^+$ T cell : DLBCL ratio. Fisher's exact test with BH-adjusted p values for comparison of categorical variables. Kruskal-Wallis test followed by post-hoc Dunn's test with adjusted p values for comparison of continuous variables.

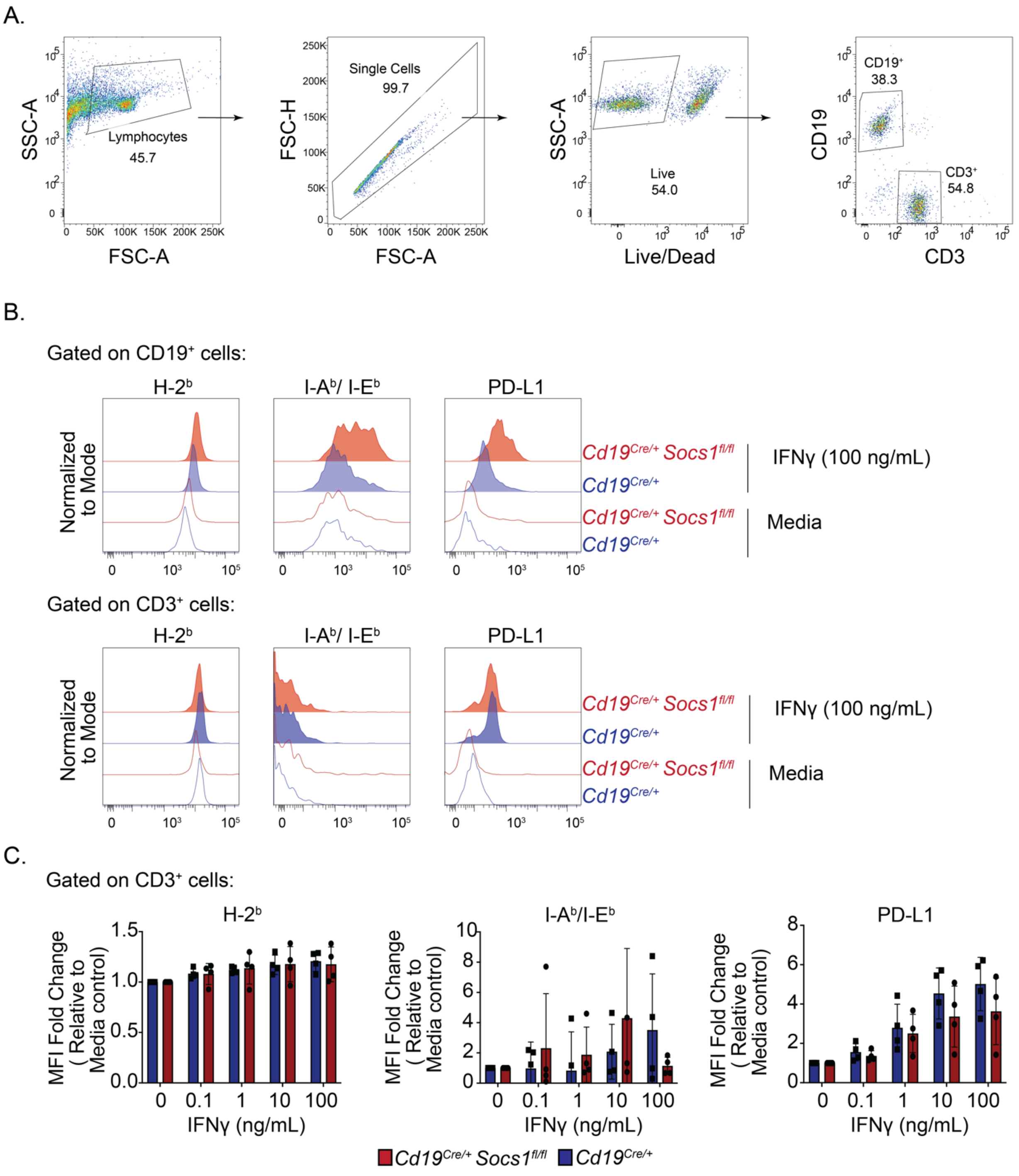


**Supplementary Figure 8. *SOCS1* mutations are enriched among GCB Hot DLBCLs and enhance B cell sensitivity to IFNγ signaling. A.** Representative gating for splenocytes isolated from *Cd19*$^{Cre/+}$ or *Cd19*$^{Cre/+}$ *Socs1*$^{fl/fl}$ mice. **B.** Representative staining of H-2$^b$, I-A/I-E$^b$, and PD-

L1 on $CD19^+$ and $CD3^+$ splenocytes from *Cd19*$^{Cre/+}$ or *Cd19*$^{Cre/+}$ *Socs1*$^{fl/fl}$ mice cultured with media or 100 ng/mL of IFNγ for 48 hours. **C.** Fold change in mean fluorescence intensity (MFI) of H-$2^b$, I-A/I-$E^b$, and PD-L1 on $CD3^+$ splenocytes. Splenocytes from *Cd19*$^{Cre/+}$ ($n = 6$) or *Cd19*$^{Cre/+}$ *Socs1*$^{fl/fl}$ mice ($n = 6$) were cultured with media or the indicated concentrations of IFNγ for 48 hours and expression levels of H-$2^b$, I-A/I-$E^b$, and PD-L1 were measured. Mice were pooled from 3 independent biological replicates. Two-way ANOVA with Bonferroni correction, adj. p values displayed (* adj. $p < 0.05$, ** adj. $p < 0.01$, *** adj. $p < 0.001$, **** adj. $p < 0.0001$).

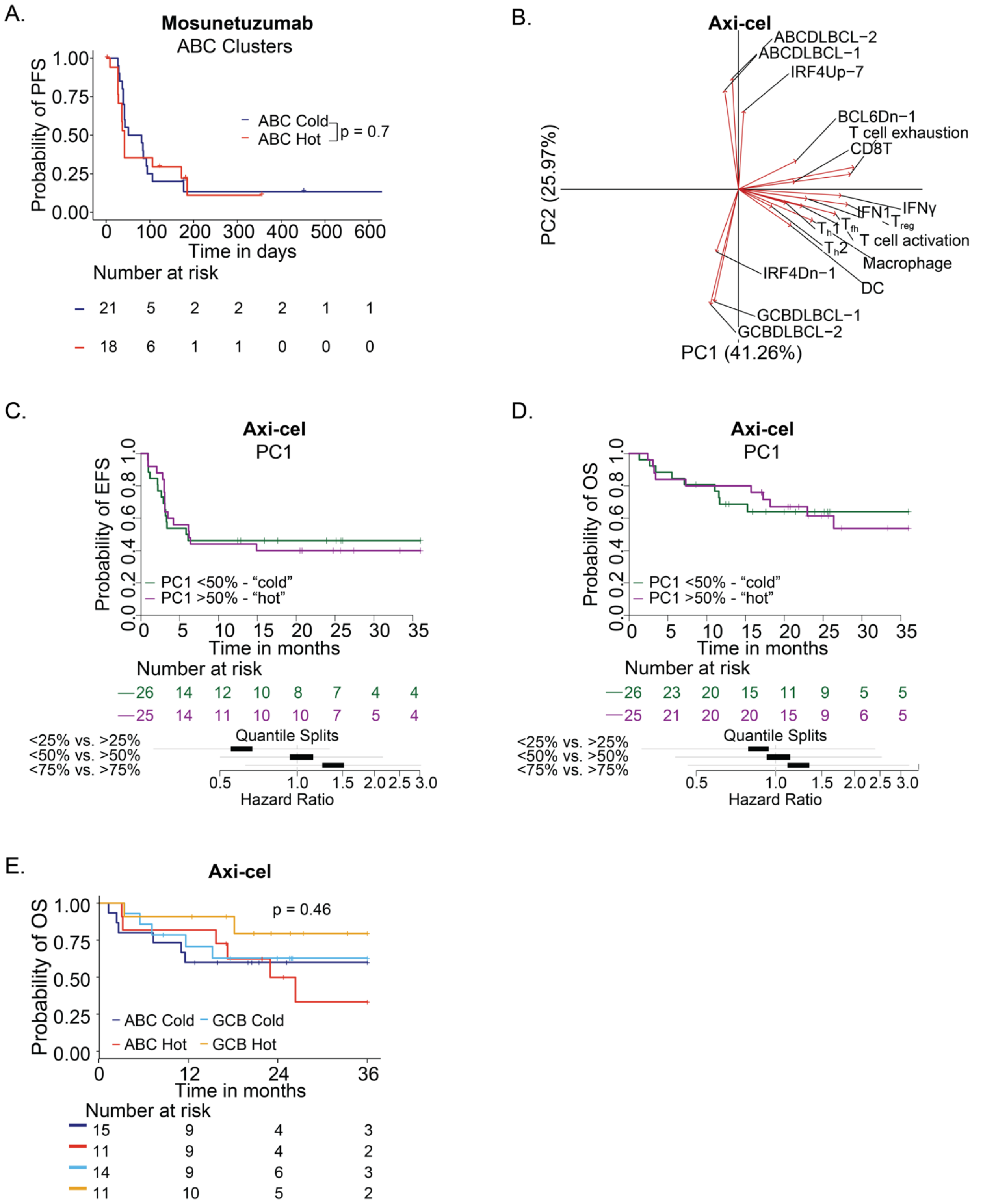


**Supplementary Figure 9. DLBCL-IQs are associated with distinct survival outcomes to BsAb therapy but not CAR T cell treatment. A.** PFS for patients assigned to ABC Hot and

ABC Cold DLBCL-IQs and treated with mosunetuzumab. Log-rank test with p value displayed. **B.** PCA biplot showing contribution of immune-related and COO-related gene sets to PC1 and PC2, respectively, for biopsies from patients treated with axi-cel. **C.** Event free survival (EFS) for patients assigned to "hot" or "cold" DLBCL immune clusters and treated with axi-cel (top) and forest plot showing hazard ratio of the association between PC1 score and EFS (bottom). **D.** OS for patients assigned to "hot" or "cold" DLBCL immune clusters and treated with axi-cel (top) and forest plot showing hazard ratio of the association between PC1 score and OS (bottom). **E.** OS for patients assigned to each DLBCL-IQ and treated with axi-cel (top) and forest plot showing hazard ratio of the association between PC1 score and OS (bottom).

**Supplementary Table 1. List of genes included in GSVA analysis.**

| Gene Set | Genes |
| --- | --- |
| **T cell exhaustion** | *LAG3 CTLA4 CD274 CD160 BTLA VSIR LAIR1 HAVCR2 CD244 TIGIT* |
| **T cell activation** | *ICOS CD28 CD27 TNFSF14 CD40LG TNFRSF9 TNFRSF4 TNFRSF25 TNFRSF18 TNFRSF8 SLAMF1 CD2 CD226* |
| **Cytolytic score** | *GZMA GZMH GZMM PRF1 GNLY* |
| **Interferon gamma** | *TIGIT CXCR6 CXCL9 CD27 CMKLR1 HLA-DQA1 CD8A NKG7 CD276 PDCD1LG2 CCL5 STAT1 LAG3 PSMB10 HLA-DRB1 CD274 IDO1 HLA-E* |
| **IFN-1** | *MX1 TNFSF10 RSAD2 IFIT1 IFIT3 IFIT2 IRF7 DDX4 MX2 ISG20* |
| **CD8T** | *ADRM1 AHSA1 C1GALT1C1 CCT6B CD37 CD3D CD3E CD3G CD69 CD8A CETN3 CSE1L GEMIN6 GNLY GPT2 GZMA GZMH GZMK IL2RB LCK MPZL1 NKG7 PIK3IP1 PTRH2 TIMM13 ZAP70* |
| **Regulatory T cell** | *FOXP3 LINC02694 IL5 CTLA4 IL32 GPR15 IL4* |
| **$T_h$2** | *ASB2 CSRP2 DAPK1 DLC1 DNAJC12 DUSP6 GNAI1 LAMP3 NRP2 OSBPL1A PDE4B PHLDA1 PLA2G4A RAB27B RBMS3 RNF125 TMPRSS3 GATA3 BIRC5 CDC25C CDC7 CENPF CXCR6 DHFR EVI5 GSTA4 HELLS IL26 LAIR2* |
| **Th1** | *CD70 TBX21 ADAM8 AHCYL2 ALCAM B3GALNT1 BBS12 BST1 CD151 CD47 CD48 CD52 CD53 CD59 CD6 CD68 CD7 CD96 CFHR3 CHRM3 CLEC7A COL23A1 COL4A4 COL5A3 DAB1 DLEU7 DOC2B EMP1 F12 FURIN GAB3 GATM GFPT2 GPR25 GREM2 HAVCR1 HSD11B1 HUNK IGF2 RCSD1 RYR1 SAV1 SELE SELP SH3KBP1 SIT1 SLC35B3 SIGLEC10 SKAP1 THUMPD2 TIGIT ZEB2 ENC1 RETREG1 FBXO30 FCGR2C STAC LTC4S MAN1B1 MDH1 MMD RGS16 IL12A P2RX5 ADGRE5 ITGB4 ICAM3 METRNL TNFRSF1A IRF1 HTR2B CALD1 MOCOS TRAF3IP2 TLR8 TRAF1 DUSP14* |
| **T follicular helper** | *B3GAT1 CDK5R1 PDCD1 BCL6 CD200 CD83 CD84 FGF2 GPR18 CEBPA ADA2 CLEC10A CLEC4A CSF1R CTSS SYNM DPP4 LRRC32 MC5R MICA NCAM1 NCR2 NRP1 PDCD1LG2 PDCD6 PRDX1 RAE1 RAET1E SIGLEC7 SIGLEC9 TYRO3 CHST12 CLIC3 IVNS1ABP KIR2DL2 LGMN* |
| **Macrophage** | *FUCA1 MMP9 LGMN HS3ST2 TM4SF19 CLEC5A GPNMB KCNJ5-AS1 CD68 CYBB* |
| **Dendritic cell** | *CD1A CD1B CD1E CCL13 CCL17 ALDH1A2 CD209 ALOX15 HLA-DQA1 FPR3* |

| | |
|---|---|
| **ABCDLBCL-1** | *ACP1 BATF BCL2 CCND2 CSNK1E ENTPD1 FUT8 GOT2 IGHG1 IL16 IRF4 MARCKS PIM1 PIM2 PRKCB PTPN1 SLA SP140 SPIB TCF4* |
| **ABCDLBCL-2** | *BLNK BMF CCDC50 CCND2 ENTPD1 ETV6 FOXP1 FUT8 IGHM IL16 IRF4 PIM1 PTPN1 SH3BP5 TBC1D27P* |
| **GCBDLBCL-1** | *BCL6 CSTB FAM3C ITPKB LMO2 IRAG2 MME MYBL1 SPINK2 VCL* |
| **GCBDLBCL-2** | *BCL6 DENND3 ITPKB LMO2 IRAG2 MME MYBL1 NEK6 SAMD12 SERPINA9* |
| **IRF4Up7** | *ALAD ANKRD33B ARHGAP17 ARHGAP24 ARHGAP25 ARHGAP31 ARHGEF3 ARID3A ASPHD2 ATP1B1 AURKA BATF BCL2 BCL3 BLNK BMF BSPRY RHEX CABLES1 CARD11 CCDC113 CCDC88C CCL22 CCND2 CD47 CDKL1 CFLAR CLINT1 COL9A2 CORO1C CSNK1E CXXC5 CYB5R2 DCTD DGKG DHRS9 DNAJC25-GNG10 DOCK10 DUSP15 DUSP5 EHD1 EHD3 EIF2S2 ELL2 ELOVL7 ENTPD1 ERP29 ETV6 FA2H FCMR FCRL5 FKBP11 FOXP1 GAB2 GID4 GYG1 HCK HIVEP2 HSP90B1 IDH1 IL10 IL16 IQGAP2 IRF2 IRF2BP2 IVNS1ABP KLHDC9 KRAS LBH LDLR LYN MAP3K5 MAPKAPK2 MARS1 MLKL MOCOS MPEG1 MSRB1 MYOCD NCF2 NDRG1 NFKBIZ NRROS OAS1 PAK2 PARVB PDCD4 PDE4B PDLIM1 PHACTR2 PIAS2 PIGR PIM1 PNP POU2F2 PPFIBP2 PRDM2 PRPF40A PTPN1 RAC2 RAPGEF1 RARA RASGRP4 RHOQ RILPL2 RUBCNL S1PR1 SACS SCD SEC11C SERPINB8 SFTPB SH2D3C SH3BP1 SH3BP5 SIDT1 SKIL SLA SLC25A30 SLC33A1 SLC39A10 SLC4A5 SMARCA2 SNX9 SP140 SPIB SPTBN1 SSR3 ST3GAL1 ST6GALNAC4 STAMBPL1 STAT3 TCF4 TCTN3 TET2 TLE1 TMEM154 TNFAIP8 TOX2 TPM4 TRAM2 TREX1 UBALD2 UCK2 UNC93B1 VASH2 VAV2 VEGFA VOPP1 WNT10A WNT9A YARS1 ZBTB32 ZFAT ZNF432* |
| **IRF4Dn1** | *ADGRG5 AIM2 ALOX5AP ANK1 ANK3 ARHGAP44 ARL5B ATL2 BCAS4 BFSP2 BICD1 BORCS8-MEF2B BPTF SHLD1 CCDC126 CCDC69 CCND3 CD1A CD27 CD38 CD83 CD86 CDK14 CEP126 CFAP58 CLIC5 COA1 COTL1 CUX1 CYP39A1 DAAM1 DEF8 DHRS9 DIP2C DOK3 EBF1 EHD3 ELF1 EML6 ENPP3 EPSTI1 ERP44 FAM53B FANCA FCRL1 FCRLB GATAD2B GCNT1 GCSAM GPR160 HECW2 HGSNAT HSD17B12 HTR3A IGSF22 ILDR1 IQCD ITPKB IZUMO4 KCNN3 GARRE1 KIAA1549L KLF12 KLHL6 KRT13 LACC1 LCK LHPP LNPEP LPP IRAG2 LY9 LYL1 MAP3K7CL MAP4K4 MAST3 MBD4 MCTP2 MED12L MEF2C MET MILR1 MOB1A MOB3A MPZL3 MTF2 MYO1E NCALD* |

| | |
|---|---|
| | *NCOA7 NLRP2 NR3C1 OTULIN PACSIN1 PAG1 PALD1 PHLPP1 PIK3CG PIP4K2A PITPNC1 PLAG1 PLXNB2 POLD4 POLH PPIL2 PRAG1 PRKCD PRKRIP1 PTAFR PTK2B PTPN18 PTPRS PUDP PXK RCBTB2 RECQL5 REL RFTN1 RRM2B S1PR2 SEC14L1 SEMA4A SGPP1 SH2B2 SH3KBP1 SLC15A4 SLC25A27 SLC2A5 SLC6A16 SMARCA4 SMIM14 SOBP SOX5 SPRED2 STAG3 STX7 SWAP70 SYK SYNE2 SYT11 TBC1D4 TMEM123 TMEM131L TMEM229B TNFSF10 TOB2 TPCN1 TPCN2 UBE2J1 USP12 VGLL4 WIPI2 XKR6 XYLT1 ZFHX3 ZNF318 ZNF581 ZNF608* |
| **BCL6Dn-1** | *ATR CCL3 CCND2 CD44 CD69 CD80 CDKN1A CDKN1B CXCL10 CXCR4 GPR183 ID2 IFITM1 IFITM3 IRF9 NFKB1 PRDM1 STAT1 TP53* |

**Supplementary Table 2. Frequency and number of DLBCL driver genes in NCI, UCMC, and BCC datasets.**

| HUGO Symbol | NCI-UCMC-BCC (%) | NCI-UCMC-BCC (n) | NCI (%) | NCI (n) | UCMC (%) | UCMC (n) | BCC (%) | BCC (n) |
|---|---|---|---|---|---|---|---|---|
| *KMT2D* | 37.26 | 313 | 34.51 | 166 | 21.05 | 16 | 46.28 | 131 |
| *PIM1* | 30.47 | 256 | 30.76 | 148 | 17.1 | 13 | 33.56 | 95 |
| *BCL2* | 27.73 | 233 | 26.19 | 126 | 13.15 | 10 | 34.27 | 97 |
| *TP53* | 25.47 | 214 | 27.23 | 131 | 19.73 | 15 | 24.02 | 68 |
| *MYD88* | 24.04 | 202 | 27.65 | 133 | 21.05 | 16 | 18.72 | 53 |
| *CREBBP* | 22.26 | 187 | 20.79 | 100 | 22.36 | 17 | 24.73 | 70 |
| *BTG2* | 21.66 | 182 | 23.07 | 111 | 9.21 | 7 | 22.61 | 64 |
| *CARD11* | 20 | 168 | 18.5 | 89 | 15.78 | 12 | 23.67 | 67 |
| *H1-4* | 20 | 168 | 19.54 | 94 | 17.1 | 13 | 21.55 | 61 |
| *TNFRSF14* | 19.76 | 166 | 18.5 | 89 | 18.42 | 14 | 22.26 | 63 |
| *TNFAIP3* | 18.8 | 158 | 25.36 | 122 | 10.52 | 8 | 9.89 | 28 |
| *B2M* | 17.97 | 151 | 17.46 | 84 | 13.15 | 10 | 20.14 | 57 |
| *BCL6* | 17.97 | 151 | 20.99 | 101 | 3.94 | 3 | 16.6 | 47 |
| *CDKN2A* | 17.85 | 150 | 27.02 | 130 | 25 | 19 | 0.35 | 1 |
| *SOCS1* | 17.73 | 149 | 15.17 | 73 | 18.42 | 14 | 21.9 | 62 |
| *TMSB4X* | 17.26 | 145 | 19.33 | 93 | 5.26 | 4 | 16.96 | 48 |
| *IRF4* | 17.02 | 143 | 15.38 | 74 | 18.42 | 14 | 19.43 | 55 |
| *KLHL6* | 17.02 | 143 | 17.46 | 84 | 6.57 | 5 | 19.08 | 54 |
| *HLA-B* | 16.9 | 142 | 24.32 | 117 | 23.68 | 18 | 2.47 | 7 |
| *TBL1XR1* | 16.54 | 139 | 20.16 | 97 | 5.26 | 4 | 13.42 | 38 |
| *BTG1* | 15.83 | 133 | 15.38 | 74 | 9.21 | 7 | 18.37 | 52 |
| *SGK1* | 15.71 | 132 | 16 | 77 | 5.26 | 4 | 18.02 | 51 |
| *CD79B* | 15.11 | 127 | 15.17 | 73 | 19.73 | 15 | 13.78 | 39 |
| *EZH2* | 15 | 126 | 13.3 | 64 | 7.89 | 6 | 19.78 | 56 |
| *CCND3* | 14.76 | 124 | 16.63 | 80 | 14.47 | 11 | 11.66 | 33 |
| *ACTB* | 14.28 | 120 | 11.64 | 56 | 13.15 | 10 | 19.08 | 54 |
| *H1-2* | 14.28 | 120 | 12.26 | 59 | 15.78 | 12 | 17.31 | 49 |
| *GNA13* | 13.8 | 116 | 9.97 | 48 | 10.52 | 8 | 21.2 | 60 |
| *KLHL14* | 13.69 | 115 | 18.5 | 89 | 9.21 | 7 | 6.71 | 19 |
| *HLA-A* | 13.21 | 111 | 17.87 | 86 | 23.68 | 18 | 2.47 | 7 |
| *ZFP36L1* | 13.09 | 110 | 11.01 | 53 | 9.21 | 7 | 17.66 | 50 |
| *CD58* | 12.85 | 108 | 15.17 | 73 | 5.26 | 4 | 10.95 | 31 |
| *MEF2B* | 12.61 | 106 | 10.18 | 49 | 7.89 | 6 | 18.02 | 51 |

| | | | | | | | | |
|---|---|---|---|---|---|---|---|---|
| *BCL11A* | 12.26 | 103 | 12.88 | 62 | 9.21 | 7 | 12.01 | 34 |
| *PRDM1* | 12.26 | 103 | 16.21 | 78 | 13.15 | 10 | 5.3 | 15 |
| *CD70* | 11.9 | 100 | 16 | 77 | 6.57 | 5 | 6.36 | 18 |
| *IRF8* | 11.78 | 99 | 12.26 | 59 | 6.57 | 5 | 12.36 | 35 |
| *TMEM30A* | 11.78 | 99 | 13.09 | 63 | 1.31 | 1 | 12.36 | 35 |
| *DTX1* | 11.3 | 95 | 17.25 | 83 | 9.21 | 7 | 1.76 | 5 |
| *MYC* | 11.07 | 93 | 9.77 | 47 | 13.15 | 10 | 12.72 | 36 |
| *ARID1A* | 10.83 | 91 | 11.43 | 55 | 3.94 | 3 | 11.66 | 33 |
| *EBF1* | 10.83 | 91 | 10.81 | 52 | NA | NA | 13.78 | 39 |
| *FOXO1* | 10.47 | 88 | 9.35 | 45 | 2.63 | 2 | 14.48 | 41 |
| *XPO1* | 10.47 | 88 | 10.6 | 51 | 17.1 | 13 | 8.48 | 24 |
| *FAS* | 10.23 | 86 | 11.43 | 55 | 5.26 | 4 | 9.54 | 27 |
| *NFKBIE* | 10.23 | 86 | 11.01 | 53 | 5.26 | 4 | 10.24 | 29 |
| *KMT2C* | 10 | 84 | 7.48 | 36 | 3.94 | 3 | 15.9 | 45 |
| *SPEN* | 10 | 84 | 12.05 | 58 | 3.94 | 3 | 8.12 | 23 |
| *EP300* | 9.64 | 81 | 9.97 | 48 | 5.26 | 4 | 10.24 | 29 |
| *PDE4DIP* | 9.52 | 80 | 12.47 | 60 | 3.94 | 3 | 6 | 17 |
| *CIITA* | 9.4 | 79 | 8.1 | 39 | 14.47 | 11 | 10.24 | 29 |
| *MCL1* | 9.28 | 78 | 9.14 | 44 | 19.73 | 15 | 6.71 | 19 |
| *STAT3* | 9.28 | 78 | 10.39 | 50 | 3.94 | 3 | 8.83 | 25 |
| *ATM* | 9.16 | 77 | 8.73 | 42 | 3.94 | 3 | 11.3 | 32 |
| *ETS1* | 9.04 | 76 | 11.85 | 57 | 3.94 | 3 | 5.65 | 16 |
| *CD83* | 8.8 | 74 | 6.65 | 32 | 10.52 | 8 | 12.01 | 34 |
| *BCL10* | 8.57 | 72 | 11.85 | 57 | 1.31 | 1 | 4.94 | 14 |
| *H1-5* | 8.57 | 72 | 11.64 | 56 | 13.15 | 10 | 2.12 | 6 |
| *SETD1B* | 8.57 | 72 | 11.85 | 57 | 11.84 | 9 | 2.12 | 6 |
| *HLA-C* | 8.45 | 71 | 10.81 | 52 | 18.42 | 14 | 1.76 | 5 |
| *ETV6* | 8.33 | 70 | 12.26 | 59 | 9.21 | 7 | 1.41 | 4 |
| *POU2F2* | 8.21 | 69 | 9.77 | 47 | 7.89 | 6 | 5.65 | 16 |
| *UBE2A* | 8.09 | 68 | 13.72 | 66 | 2.63 | 2 | NA | NA |
| *DUSP2* | 7.85 | 66 | 12.26 | 59 | 7.89 | 6 | 0.35 | 1 |
| *H2BC5* | 7.85 | 66 | 11.01 | 53 | 9.21 | 7 | 2.12 | 6 |
| *LTB* | 7.85 | 66 | 10.18 | 49 | 13.15 | 10 | 2.47 | 7 |
| *ATR* | 7.73 | 65 | 9.97 | 48 | 2.63 | 2 | 5.3 | 15 |
| *BIRC6* | 7.73 | 65 | 6.44 | 31 | 6.57 | 5 | 10.24 | 29 |
| *DDX3X* | 7.73 | 65 | 9.35 | 45 | 1.31 | 1 | 6.71 | 19 |
| *GNAI2* | 7.5 | 63 | 10.81 | 52 | 3.94 | 3 | 2.82 | 8 |
| *ZC3H12A* | 7.5 | 63 | 7.27 | 35 | 15.78 | 12 | 5.65 | 16 |
| *CD274* | 7.26 | 61 | 8.73 | 42 | 10.52 | 8 | 3.88 | 11 |
| *H2AC17* | 7.14 | 60 | 9.56 | 46 | 10.52 | 8 | 2.12 | 6 |

| | | | | | | | | |
|---|---|---|---|---|---|---|---|---|
| *ARID1B* | 7.02 | 59 | 11.22 | 54 | 3.94 | 3 | 0.7 | 2 |
| *BRAF* | 7.02 | 59 | 7.48 | 36 | NA | NA | 8.12 | 23 |
| *CHST2* | 7.02 | 59 | 8.31 | 40 | 3.94 | 3 | 5.65 | 16 |
| *PAX5* | 7.02 | 59 | 10.81 | 52 | 6.57 | 5 | 0.7 | 2 |
| *IL6* | 6.9 | 58 | 6.02 | 29 | 5.26 | 4 | 8.83 | 25 |
| *POU2AF1* | 6.9 | 58 | 7.48 | 36 | 9.21 | 7 | 5.3 | 15 |
| *RHOA* | 6.9 | 58 | 7.9 | 38 | 2.63 | 2 | 6.36 | 18 |
| *TOX* | 6.9 | 58 | 11.01 | 53 | 2.63 | 2 | 1.06 | 3 |
| *H1-3* | 6.78 | 57 | 9.14 | 44 | 10.52 | 8 | 1.76 | 5 |
| *H2AC6* | 6.78 | 57 | 9.14 | 44 | 9.21 | 7 | 2.12 | 6 |
| *NAV1* | 6.78 | 57 | 6.44 | 31 | 3.94 | 3 | 8.12 | 23 |
| *NOTCH2* | 6.66 | 56 | 9.97 | 48 | 9.21 | 7 | 0.35 | 1 |
| *CBLB* | 6.42 | 54 | 7.9 | 38 | 2.63 | 2 | 4.94 | 14 |
| *FOXP1* | 6.42 | 54 | 7.48 | 36 | 7.89 | 6 | 4.24 | 12 |
| *MECOM* | 6.42 | 54 | 7.48 | 36 | 3.94 | 3 | 5.3 | 15 |
| *PIM2* | 6.42 | 54 | 10.81 | 52 | 2.63 | 2 | NA | NA |
| *RB1* | 6.42 | 54 | 5.61 | 27 | NA | NA | 9.54 | 27 |
| *SETD2* | 6.42 | 54 | 7.48 | 36 | 1.31 | 1 | 6 | 17 |
| *H2BC21* | 6.3 | 53 | 6.23 | 30 | 9.21 | 7 | 5.65 | 16 |
| *STAT6* | 6.19 | 52 | 5.82 | 28 | 7.89 | 6 | 6.36 | 18 |
| *CDC73* | 6.07 | 51 | 6.02 | 29 | 6.57 | 5 | 6 | 17 |
| *NFKBIA* | 6.07 | 51 | 5.19 | 25 | 6.57 | 5 | 7.42 | 21 |
| *BCL7A* | 5.83 | 49 | 6.86 | 33 | 9.21 | 7 | 3.18 | 9 |
| *DCAF6* | 5.83 | 49 | 5.19 | 25 | 9.21 | 7 | 6 | 17 |
| *H2BC12* | 5.83 | 49 | 7.27 | 35 | 10.52 | 8 | 2.12 | 6 |
| *TIPARP* | 5.83 | 49 | 7.27 | 35 | 2.63 | 2 | 4.24 | 12 |
| *HLA-DMA* | 5.71 | 48 | 5.82 | 28 | 17.1 | 13 | 2.47 | 7 |
| *MGA* | 5.71 | 48 | 9.77 | 47 | 1.31 | 1 | NA | NA |
| *NCOR1* | 5.71 | 48 | 8.31 | 40 | 6.57 | 5 | 1.06 | 3 |
| *BRINP3* | 5.59 | 47 | 5.19 | 25 | 6.57 | 5 | 6 | 17 |
| *MET* | 5.59 | 47 | 6.02 | 29 | 1.31 | 1 | 6 | 17 |
| *TET2* | 5.47 | 46 | 8.31 | 40 | 5.26 | 4 | 0.7 | 2 |
| *KRAS* | 5.35 | 45 | 5.61 | 27 | 2.63 | 2 | 5.65 | 16 |
| *SETD5* | 5.23 | 44 | 7.06 | 34 | 2.63 | 2 | 2.82 | 8 |
| *ZNF292* | 5.23 | 44 | 8.52 | 41 | 2.63 | 2 | 0.35 | 1 |
| *BTK* | 5.11 | 43 | 8.31 | 40 | 3.94 | 3 | NA | NA |
| *H3C2* | 5.11 | 43 | 3.95 | 19 | 10.52 | 8 | 5.65 | 16 |
| *HNRNPU* | 5.11 | 43 | 6.65 | 32 | 2.63 | 2 | 3.18 | 9 |
| *NLRP8* | 5 | 42 | 5.19 | 25 | 9.21 | 7 | 3.53 | 10 |
| *SIN3A* | 5 | 42 | 5.82 | 28 | 1.31 | 1 | 4.59 | 13 |

| | | | | | | | | |
|---|---|---|---|---|---|---|---|---|
| *PTPN6* | 4.88 | 41 | 6.23 | 30 | 7.89 | 6 | 1.76 | 5 |
| *S1PR2* | 4.88 | 41 | 4.15 | 20 | 7.89 | 6 | 5.3 | 15 |
| *DDX10* | 4.76 | 40 | 6.02 | 29 | 1.31 | 1 | 3.53 | 10 |
| *EEF1A1* | 4.76 | 40 | 7.48 | 36 | 1.31 | 1 | 1.06 | 3 |
| *INO80* | 4.76 | 40 | 7.48 | 36 | 5.26 | 4 | NA | NA |
| *MARK1* | 4.64 | 39 | 5.19 | 25 | 3.94 | 3 | 3.88 | 11 |
| *TAF1* | 4.64 | 39 | 7.69 | 37 | 2.63 | 2 | NA | NA |
| *ZEB2* | 4.64 | 39 | 7.06 | 34 | 3.94 | 3 | 0.7 | 2 |
| *ZFAT* | 4.64 | 39 | 4.78 | 23 | 6.57 | 5 | 3.88 | 11 |
| *CRIP1* | 4.52 | 38 | 3.11 | 15 | 9.21 | 7 | 5.65 | 16 |
| *ZNF608* | 4.52 | 38 | 6.65 | 32 | 2.63 | 2 | 1.41 | 4 |
| *IKZF3* | 4.28 | 36 | 5.61 | 27 | 2.63 | 2 | 2.47 | 7 |
| *MAGT1* | 4.28 | 36 | 7.06 | 34 | 2.63 | 2 | NA | NA |
| *MYB* | 4.28 | 36 | 7.06 | 34 | 2.63 | 2 | NA | NA |
| *PHF6* | 4.28 | 36 | 6.23 | 30 | 1.31 | 1 | 1.76 | 5 |
| *PRKCB* | 4.16 | 35 | 6.02 | 29 | 2.63 | 2 | 1.41 | 4 |
| *PTPRK* | 4.16 | 35 | 7.06 | 34 | 1.31 | 1 | NA | NA |
| *TGFBR2* | 4.16 | 35 | 4.57 | 22 | 1.31 | 1 | 4.24 | 12 |
| *WAC* | 4.16 | 35 | 5.61 | 27 | 5.26 | 4 | 1.41 | 4 |
| *ZFX* | 4.16 | 35 | 7.06 | 34 | 1.31 | 1 | NA | NA |
| *IL16* | 4.04 | 34 | 6.23 | 30 | 3.94 | 3 | 0.35 | 1 |
| *MSH6* | 4.04 | 34 | 4.57 | 22 | 2.63 | 2 | 3.53 | 10 |
| *PIK3CD* | 4.04 | 34 | 6.44 | 31 | 1.31 | 1 | 0.7 | 2 |
| *PTEN* | 4.04 | 34 | 6.86 | 33 | 1.31 | 1 | NA | NA |
| *SMARCA4* | 4.04 | 34 | 3.74 | 18 | 7.89 | 6 | 3.53 | 10 |
| *MSH2* | 3.92 | 33 | 4.36 | 21 | 2.63 | 2 | 3.53 | 10 |
| *SYK* | 3.92 | 33 | 4.78 | 23 | 2.63 | 2 | 2.82 | 8 |
| *RARA* | 3.8 | 32 | 4.57 | 22 | 2.63 | 2 | 2.82 | 8 |
| *UBR5* | 3.8 | 32 | 5.4 | 26 | 2.63 | 2 | 1.41 | 4 |
| *MAP2K1* | 3.69 | 31 | 6.23 | 30 | 1.31 | 1 | NA | NA |
| *CHD8* | 3.45 | 29 | 3.32 | 16 | NA | NA | 4.59 | 13 |
| *HVCN1* | 3.45 | 29 | 3.32 | 16 | 3.94 | 3 | 3.53 | 10 |
| *IKBKB* | 3.45 | 29 | 5.19 | 25 | 1.31 | 1 | 1.06 | 3 |
| *LYN* | 3.45 | 29 | 4.78 | 23 | 3.94 | 3 | 1.06 | 3 |
| *RUNX1* | 3.45 | 29 | 3.53 | 17 | 6.57 | 5 | 2.47 | 7 |
| *GRB2* | 3.33 | 28 | 4.36 | 21 | 2.63 | 2 | 1.76 | 5 |
| *HRAS* | 3.33 | 28 | 2.91 | 14 | 11.84 | 9 | 1.76 | 5 |
| *PRPS1* | 3.33 | 28 | 5.61 | 27 | 1.31 | 1 | NA | NA |
| *YY1* | 3.33 | 28 | 4.78 | 23 | 6.57 | 5 | NA | NA |
| *MTOR* | 3.21 | 27 | 5.4 | 26 | NA | NA | 0.35 | 1 |

| | | | | | | | | |
|---|---|---|---|---|---|---|---|---|
| *DNMT3A* | 3.09 | 26 | 3.11 | 15 | 6.57 | 5 | 2.12 | 6 |
| *JUNB* | 3.09 | 26 | 3.32 | 16 | 10.52 | 8 | 0.7 | 2 |
| *TLR2* | 2.97 | 25 | 4.57 | 22 | NA | NA | 1.06 | 3 |
| *FBXW7* | 2.85 | 24 | 4.36 | 21 | NA | NA | 1.06 | 3 |
| *RAD9A* | 2.85 | 24 | 2.7 | 13 | 6.57 | 5 | 2.12 | 6 |
| *ANKRD17* | 2.73 | 23 | 4.36 | 21 | 1.31 | 1 | 0.35 | 1 |
| *ZBTB7A* | 2.73 | 23 | 2.07 | 10 | 9.21 | 7 | 2.12 | 6 |
| *ARID5B* | 2.61 | 22 | 4.36 | 21 | NA | NA | 0.35 | 1 |
| *CD22* | 2.61 | 22 | 3.32 | 16 | 2.63 | 2 | 1.41 | 4 |
| *NF1* | 2.61 | 22 | 3.11 | 15 | 2.63 | 2 | 1.76 | 5 |
| *IGLL5* | 2.38 | 20 | 1.03 | 5 | 17.1 | 13 | 0.7 | 2 |
| *KCMF1* | 2.38 | 20 | 3.32 | 16 | 2.63 | 2 | 0.7 | 2 |
| *PIK3R1* | 2.38 | 20 | 1.24 | 6 | 2.63 | 2 | 4.24 | 12 |
| *RRAGC* | 2.38 | 20 | 3.95 | 19 | 1.31 | 1 | NA | NA |
| *HNRNPD* | 2.26 | 19 | 3.74 | 18 | NA | NA | 0.35 | 1 |
| *JAK1* | 2.26 | 19 | 3.74 | 18 | NA | NA | 0.35 | 1 |
| *SF3B1* | 2.26 | 19 | 3.11 | 15 | 1.31 | 1 | 1.06 | 3 |
| *CXCR4* | 2.14 | 18 | 2.7 | 13 | 6.57 | 5 | NA | NA |
| *NFKB2* | 2.14 | 18 | 3.32 | 16 | 1.31 | 1 | 0.35 | 1 |
| *TCL1A* | 2.14 | 18 | 3.53 | 17 | NA | NA | 0.35 | 1 |
| *JAK3* | 2.02 | 17 | 1.24 | 6 | 10.52 | 8 | 1.06 | 3 |
| *CCL4* | 1.9 | 16 | 1.45 | 7 | 3.94 | 3 | 2.12 | 6 |
| *CHD1* | 1.9 | 16 | 2.7 | 13 | NA | NA | 1.06 | 3 |
| *FUT5* | 1.9 | 16 | 1.24 | 6 | 6.57 | 5 | 1.76 | 5 |
| *COQ7* | 1.78 | 15 | 1.87 | 9 | 1.31 | 1 | 1.76 | 5 |
| *GNAS* | 1.78 | 15 | 1.24 | 6 | 6.57 | 5 | 1.41 | 4 |
| *BTBD3* | 1.66 | 14 | 1.45 | 7 | 1.31 | 1 | 2.12 | 6 |
| *NANOG* | 1.66 | 14 | 1.24 | 6 | 3.94 | 3 | 1.76 | 5 |
| *PPP4R3A* | 1.66 | 14 | 2.49 | 12 | 1.31 | 1 | 0.35 | 1 |
| *CASP8* | 1.54 | 13 | 1.45 | 7 | 3.94 | 3 | 1.06 | 3 |
| *MAP4K4* | 1.54 | 13 | 2.07 | 10 | 2.63 | 2 | 0.35 | 1 |
| *STAT5B* | 1.54 | 13 | 0.83 | 4 | 2.63 | 2 | 2.47 | 7 |
| *LIN54* | 1.07 | 9 | 1.66 | 8 | NA | NA | 0.35 | 1 |
| *ZNF423* | 1.07 | 9 | 1.66 | 8 | NA | NA | 0.35 | 1 |
| *DICER1* | 0.95 | 8 | 1.45 | 7 | 1.31 | 1 | NA | NA |
| *FUBP1* | 0.95 | 8 | 1.45 | 7 | 1.31 | 1 | NA | NA |
| *GOLGA5* | 0.83 | 7 | 1.45 | 7 | NA | NA | NA | NA |
